# Single-cell spatial multiomics identifies *POSTN*^+^ CAFs mediating chemoradiotherapy resistance in rectal cancer

**DOI:** 10.64898/2026.04.30.721803

**Authors:** Shunsuke A Sakai, Masayuki Okumura, Yukiko Morinaga, Ken Kato, Motohiro Kojima, Falko Hofmann, Brian Reichholf, Pablo Vieyra Garcia, Yuka Nakamura, Shingo Sakashita, Masaki Nakamura, Hidehiro Hojo, Yuichiro Tsukada, Masaaki Ito, Kohei Shitara, Hideaki Bando, Takashi Kojima, Junko Zenkoh, Shotaro Tatekawa, Shohei Katsuki, Kazuhiko Ogawa, Yutaka Takahashi, Ayako Suzuki, Yutaka Suzuki, Katsuya Tsuchihara, Gabriela Gremel, Riu Yamashita, Shun-Ichiro Kageyama

**Author notes:** **Corresponding author:** Shun-Ichiro Kageyama, Division of Translational Informatics, Exploratory Oncology Research & Clinical Trial Center, National Cancer Center; Kashiwa, Chiba, 277-8577, Japan,; Riu Yamashita, Division of Translational Informatics, Exploratory Oncology Research & Clinical Trial Center, National Cancer Center; Kashiwa, Chiba, 277-8577, Japan.

## Abstract

Neoadjuvant chemoradiotherapy (CRT) is standard for locally advanced rectal cancer (LARC), yet many patients retain residual disease. To resolve CRT-associated remodeling of the tumor microenvironment, we generated a multimodal spatial atlas from serial sections of paired pretreatment and post-treatment specimens from 24 patients using Xenium single-cell spatial transcriptomics and PhenoCycler multiplex proteomics, profiling 2.8 million cells; matched Visium HD datasets were generated on adjacent serial sections. Resistance was most strongly associated with fibroblast and myeloid programs adjacent to residual tumor. We identify a periostin (*POSTN*)-expressing CAF subset selectively enriched around residual tumor cells in non-responders, displaying a myofibroblastic phenotype and activating extracellular matrix remodeling, noncanonical WNT signaling, and immunosuppressive pathways. Tumor cells neighboring *POSTN*^+^ CAFs show consistent epithelial–mesenchymal transition signatures. Together, this atlas enables interrogation of CRT-induced spatial remodeling and nominates *POSTN*^+^ CAFs as key mediators and targets of CRT resistance, with direct relevance to CRT-based combination strategies.

## Introduction

Colorectal cancer (CRC) is among the most prevalent malignancies globally, with approximately 700,000 new cases of rectal cancer reported annually^1^. Locally advanced rectal cancer (LARC), characterized by tumors invading beyond the rectal wall or involving regional lymph nodes, presents significant therapeutic challenges due to its high recurrence rates^2, 3^. The current standard of care for LARC includes neoadjuvant chemoradiotherapy (CRT) followed by surgical resection^4^. While CRT primarily aims to improve local control, recent evidence indicates that some patients achieve complete tumor regression (a pathological complete response, pCR) with CRT alone, potentially eliminating the need for surgery^5, 6, 7^. Avoiding surgery is particularly important for quality of life, as rectal cancer resection typically results in the need for a permanent colostomy^8^; patients achieving a pCR exhibit significantly lower recurrence risks post-surgery; however, approximately two-thirds of patients do not achieve a pCR, instead showing residual tumor cells categorized as a major pathological response (MPR) or nonresponse (NR) characterized by minimal therapeutic effect and a substantial number of residual tumor cells^4^. These patients with residual tumors face substantially higher recurrence risks, negatively affecting their survival and quality of life^9^. Consequently, understanding and overcoming CRT resistance remains a critical clinical priority in CRC treatment.

To enhance therapeutic efficacy, intensified CRT regimens, such as total neoadjuvant therapy (TNT), escalated radiation doses, and integration of immune checkpoint inhibitors, have been developed^10, 11, 12, 13, 14, 15^. These intensified treatments primarily target patients at high risk of insufficient response to standard CRT. Although these regimens have led to promising outcomes in selected patient populations, they do not uniformly improve overall survival across all patients and are often associated with increased toxicities. Thus, the novel treatment strategies being developed based on CRT are not yet satisfactory, making it crucial to elucidate the mechanisms underlying CRT resistance and identify specific and clinically valuable therapeutic targets. The mechanisms underlying CRT resistance in LARC are multifaceted and incompletely understood. Radiotherapy (RT), a fundamental component of CRT, primarily induces tumor cell death by causing severe DNA damage, particularly double-strand breaks^16, 17^. Factors such as DNA repair capacity of cancer cells and tumor hypoxia have been known to influence treatment efficacy^18^. Additionally, RT is increasingly recognized for its ability to stimulate immune responses, significantly reshaping the tumor microenvironment (TME). TME components, including M2 macrophages^19, 20, 21^, regulatory T cells (Tregs)^22, 23^, and cancer-associated fibroblasts (CAFs)^24, 25, 26^, have been implicated in treatment resistance. Specifically, CAFs contribute to radioresistance through the production of soluble factors, including IL-6 and CXCL1. However, definitive conclusions regarding the critical cell types and molecular mechanisms remain elusive. This ambiguity may arise because pivotal TME factors vary widely among cancer types, and their interactions are highly complex and spatially organized. Therefore, comprehensive spatial omics analysis that preserves native tissue architecture is essential to unravel these intricate cellular interactions. This approach will elucidate the key mechanisms underlying TME-mediated treatment resistance by capturing the dynamic spatial reorganization of cellular components following CRT.

In this study, we aimed to elucidate the cellular and molecular mechanisms underlying CRT resistance in patients with LARC by comprehensively analyzing TME. Utilizing advanced single-cell spatial transcriptomics (Xenium 5K) combined with multiplex immunostaining (PhenoCycler), we comprehensively analyzed paired pre-and post-treatment samples from 24 LARC patients (48 paired samples in total) and obtained four early post-treatment biopsy samples (JustAfter). This state-of-the-art methodology enables simultaneous single-cell-level analysis of previously reported resistance factors^27, 28^, including tumor cells, various TME components, and hypoxic conditions. This approach allowed us to identify dominant cellular and genetic contributors to treatment resistance. Our findings address critical knowledge gaps and provide a solid foundation for developing targeted therapies to effectively overcome CRT resistance, ultimately aiming to improve patient outcomes.

## Results

### Spatiotemporal multiomics profiling of patients with LARC

To delineate the spatial and molecular dynamics of LARC in response to neoadjuvant CRT, we analyzed 52 tissue samples—including 28 biopsy samples (24 pretreatment and 4 JustAfter) and 24 posttreatment resected tumor samples—from 24 LARC patients (Fig. 1a). Sampling was conducted across three clinically relevant timepoints: “Pre” CRT, “JustAfter” CRT (within 2 weeks post-CRT), and “Resection” (approximately 3 months post-CRT). A total of 52 samples were collected from 24 patients, comprising 24 samples at the “Pre” timepoint, four at “JustAfter” (two MPR and two NR), and 24 at the “Resection” timepoint. Patients were stratified into three groups on the basis of the histopathological response assessment of their resected specimens: pathological complete response (pCR, n = 8; no residual viable tumor cells), major pathological response (MPR, n = 8; less than 10% residual viable tumor cells), and no response (NR, n = 8; 10% or more residual viable tumor cells) (Table 1). To capture the spatial architecture and cellular heterogeneity of the TME, we employed a multimodal spatial omics pipeline integrating Xenium-based transcriptomics, PhenoCycler-based proteomics, and whole-exome sequencing (WES) of morphologically matched serial sections (Fig. 1b). Patient characteristics and modality availability are summarized in Fig. 1c. This integrated dataset enabled us to characterize the tumor microenvironment after chemoradiotherapy, screen for cell types associated with treatment resistance, and investigate resistance mechanisms in cancer cells. In addition, we generated transcriptome-wide spatial gene expression profiles using Visium HD (10x Genomics) on adjacent serial sections as a complementary dataset for future integrative analyses; Visium HD data were not used for the primary analyses presented in this study.

**Fig. 1.**
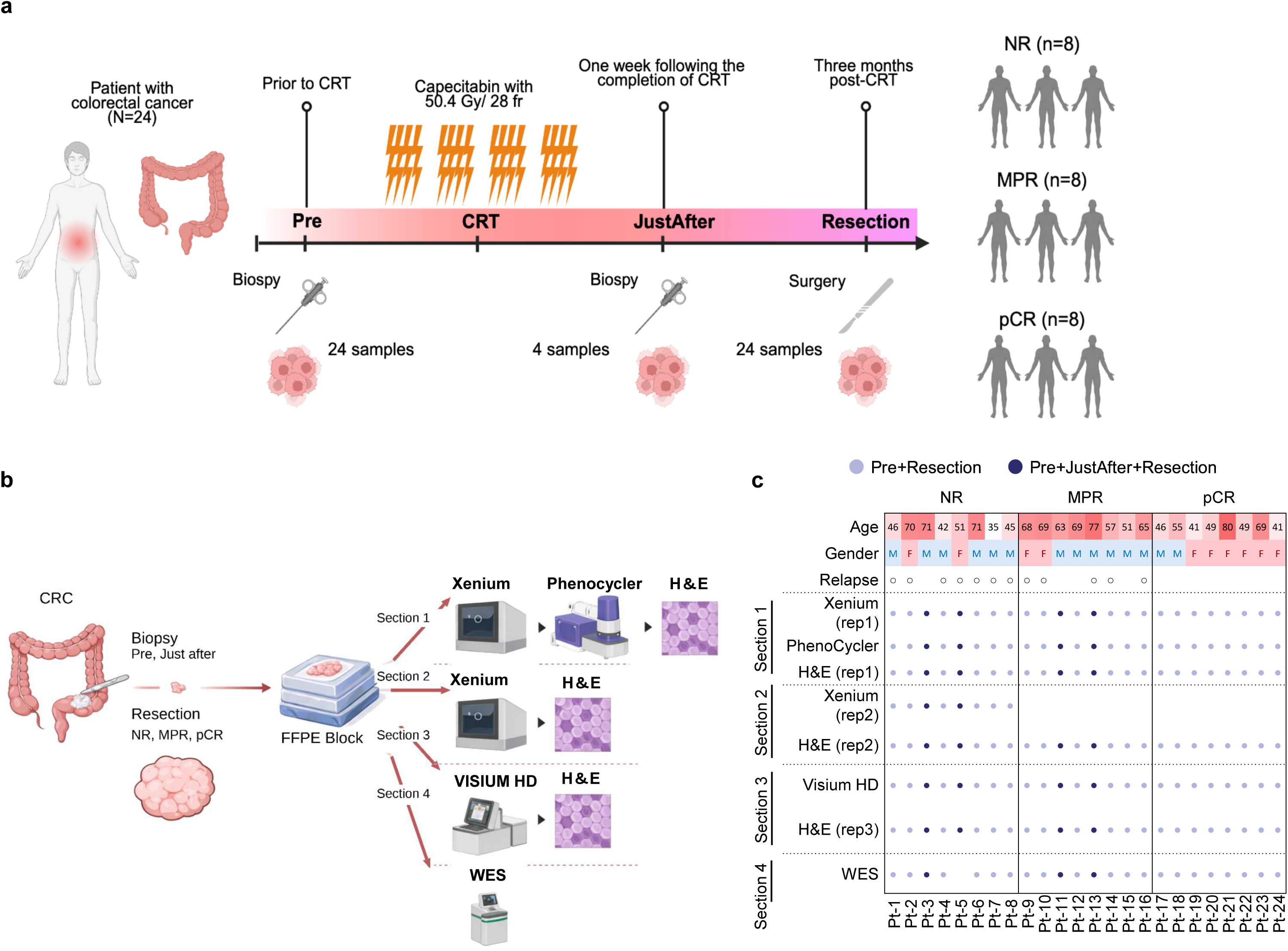
Overview of the study design, sampling time points, and multimodal dataset composition. a, Schematic illustration of the neoadjuvant chemoradiotherapy (CRT) schedule and timing of tumor sampling in the study cohort (N = 24). The CRT regimen consisted of capecitabine in combination with 50.4 Gy of radiation delivered in 28 fractions. “Pre” refers to the time point prior to CRT, “JustAfter” indicates within 2 weeks after the completion of CRT, and “Resection” represents three months post-CRT. Biopsy samples were collected from all 24 patients at the Pre time point and from 4 patients at the JustAfter time point, and surgical samples were obtained from all 24 patients at the Resection time point. Among the 24 patients, 8 achieved a pathological complete response (pCR), 8 achieved a major pathological response (MPR), and 8 were classified as no response (NR). b, Each sample obtained at the designated time points was serially sectioned into four parts (Sections 1–4). Section 1 was used for Xenium analysis, PhenoCycler analysis, and hematoxylin and eosin (H&E) staining. Section 2 was used for Xenium analysis and H&E staining. Section 3 was used for Visium HD spatial transcriptomics on adjacent serial sections. Section 4 was used for whole-exome sequencing. c, Summary matrix showing patient characteristics (age, sex, recurrence status) and the availability of each modality. The purple circles represent samples analyzed at both the Pre and Resection time points, whereas the dark circles indicate samples analyzed at all three time points: Pre, JustAfter, and Resection.

**Table 1.**
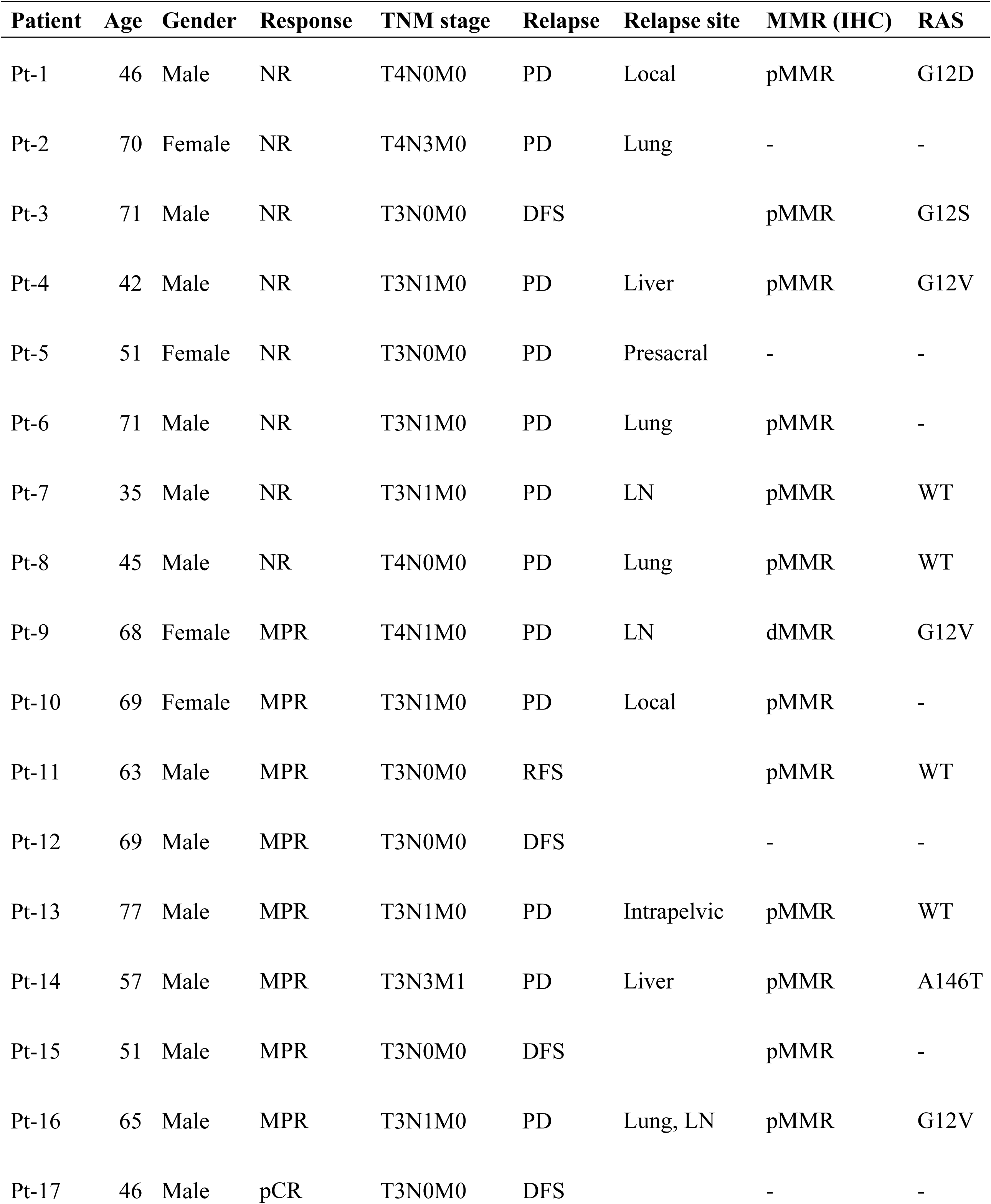
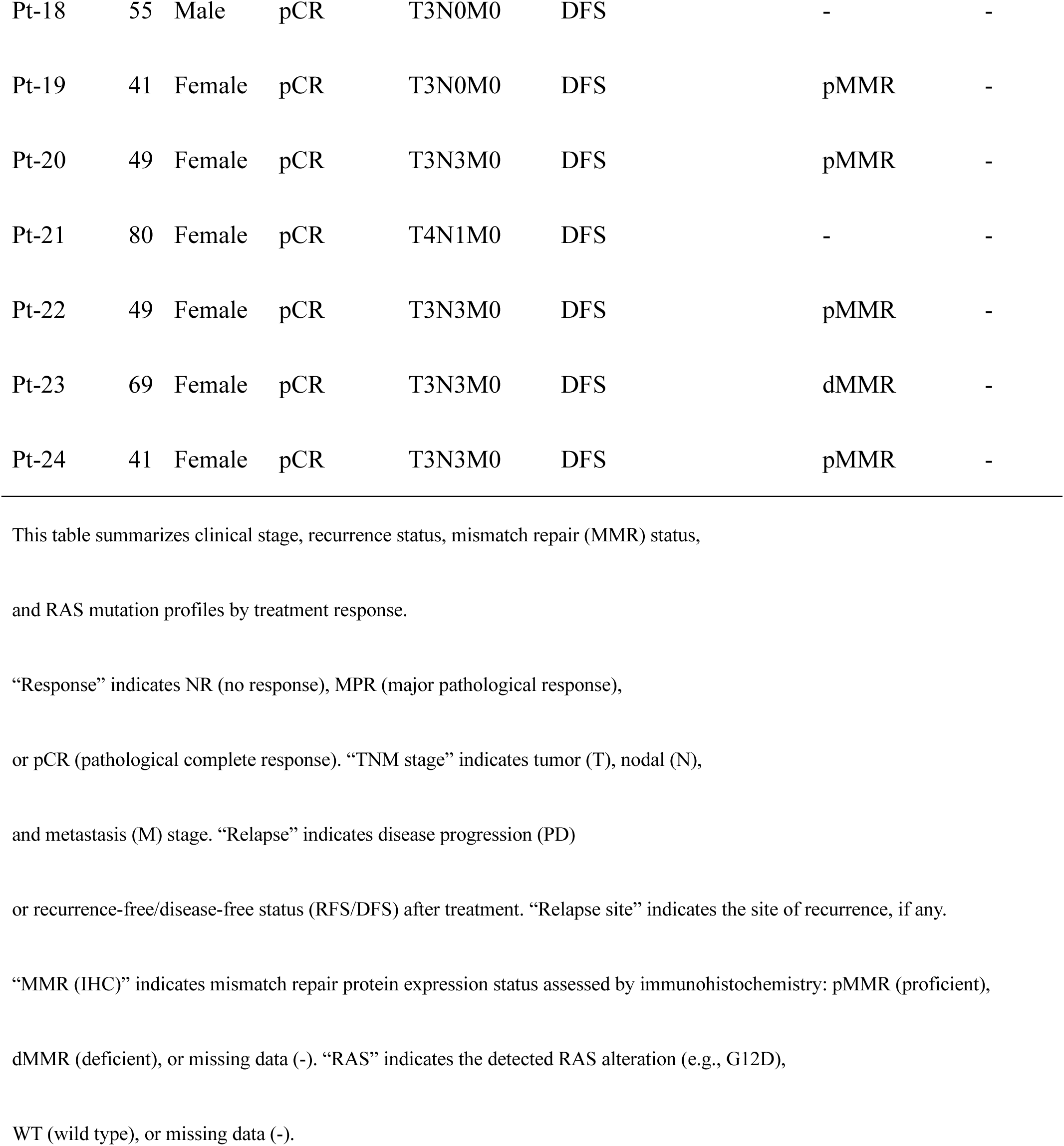
Clinical and Molecular Characteristics of Patients Based on Treatment Response.

### Spatial transcriptomics reveals dynamic changes in cellular composition pre- and post-chemoradiotherapy

To systematically characterize the cellular composition and spatial organization of the TME in LARC patients, we performed cell type identification using Xenium spatial transcriptomics data. Following stringent quality control (see Methods), we obtained 2.83 million high-quality cells, which were categorized into epithelial (39%), stromal (38%), T (9.9%), myeloid (6.0%), and B (7.5%) cell populations (Fig. 2a). The accuracy of these annotations was assessed by examining marker gene expression patterns within uniform manifold approximation and projection (UMAP) embeddings and confirmed through visual inspection of cluster-specific expression profiles (Supplementary Fig. 1a). Tumor and normal epithelial cells were spatially delineated by three pathologists who examined matched hematoxylin and eosin (H&E)-stained sections (Fig. 2b and Supplementary Fig. 1b). While conventional single-cell RNA sequencing made it difficult to distinguish between normal epithelium and tumor regions, our spatial transcriptomic dataset, combined with pathologist-guided annotation, enabled accurate distinction of normal and cancerous tissues, allowing for focused analysis of residual cancer cells (Fig. 2c). The residual tumor cells in the NR and MPR groups retained their malignant morphologies, and the pCR group samples exhibited lymphocyte-rich tumor beds, a characteristic consistent with complete regression^29^. These findings underscore the importance of spatial analysis focused on the tumor regions, where TME components are predominantly enriched. Moreover, comparisons across timepoints and response groups revealed changes in cellular composition (Fig. 2d). Specifically, the proportion of tumor cells was significantly decreased in pCR versus NR samples (*P* < 0.001) (Fig. 2e), while the proportion of stromal cells significantly increased from Pre to Resection timepoints (*P* < 0.001) (Fig. 2f). Thus, our Xenium dataset accurately captured key pathological features across response groups and timepoints throughout CRT.

**Fig. 2.**
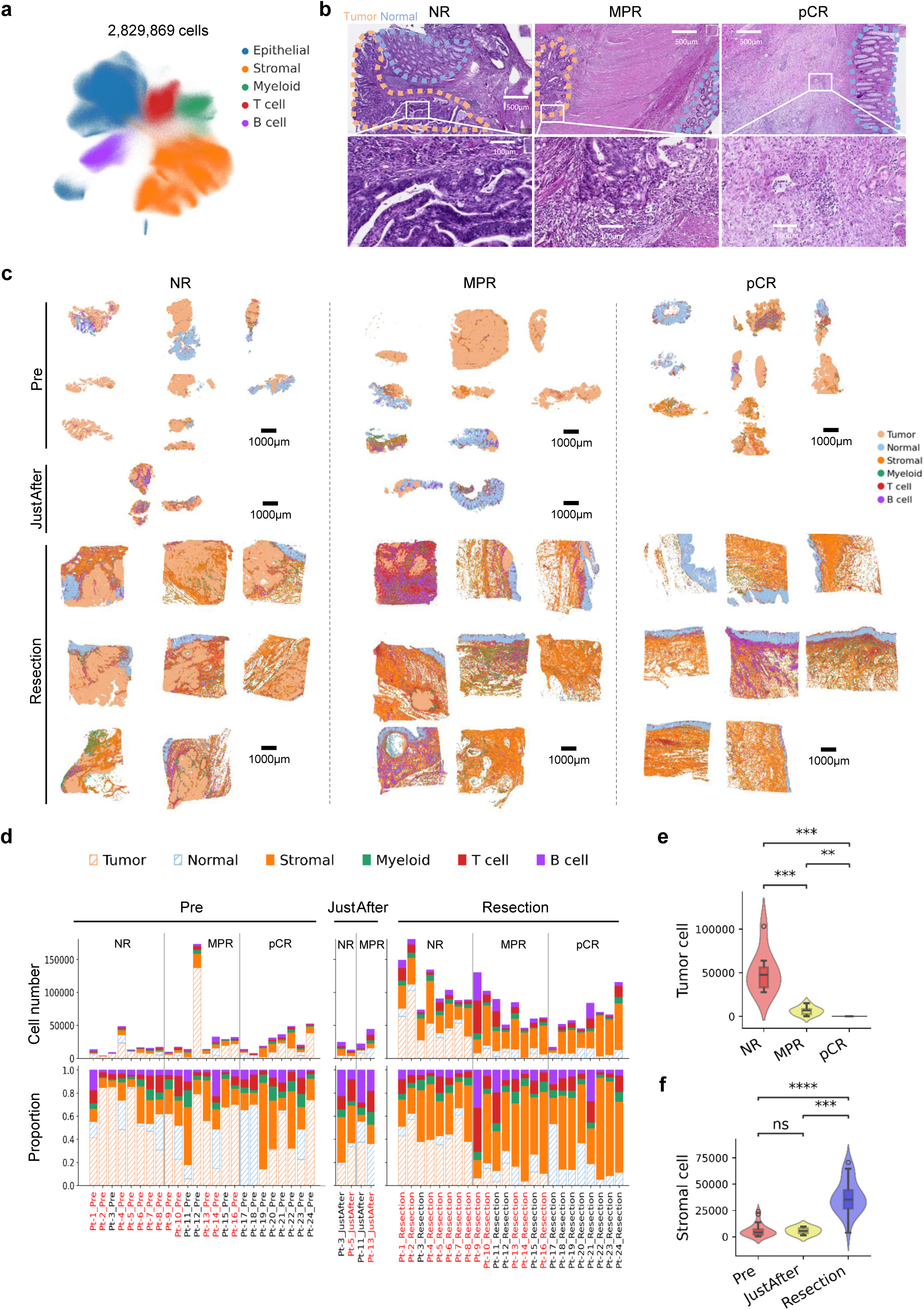
Clustering and annotation of spatial transcriptomic cell populations. a, Uniform manifold approximation and projection (UMAP) embedding of gene expression profiles from 2,829,869 cells across 52 samples from 24 patients, based on the integrated Xenium dataset. Blue indicates epithelial cells, orange indicates stromal cells, green represents myeloid cells, red corresponds to T cells, and purple denotes B cells. b, Hematoxylin and eosin (H&E)-stained tissue sections from patients in the NR, MPR, and pCR groups. The upper panels show low-magnification images; the lower panels show high-magnification images. The tumor regions are outlined with light orange wavy lines, and the normal epithelial regions are outlined with light blue wavy lines. c, Spatial localization of major clusters across all 52 samples from 24 patients, stratified by both treatment response (NR, MPR, pCR) and sampling timing (Pre, JustAfter, Resection). d, Stacked bar plots showing the total number of cells (top) and cell composition (bottom) for each sample, stratified by time point and treatment response. On the x-axis, black text indicates samples from patients without recurrence, and red indicates those from patients who experienced recurrence. e, Violin plots showing tumor cell counts stratified by the pathological response. Statistical significance was assessed using the Mann-Whitney U test. *, **, and *** indicate P < 0.05, P < 0.01, and P < 0.001, respectively. f, Violin plots showing stromal cell counts stratified by timepoint.

To evaluate interpatient variability in gene expression within major cell types, we performed dimensionality reduction analysis. Notably, tumor cells exhibited strong patient-specific clustering, in contrast to normal epithelial cells and other TME cell types, which showed less interpatient variability (Supplementary Fig. 1c). To assess whether tumor cell heterogeneity reflected major driver alterations, we analyzed whole-exome sequencing data but found no clear associations (Supplementary Fig. 1d). Similarly, mismatch repair-deficient (dMMR) tumors, as assessed by immunohistochemistry (IHC), showed no significant differences in clustering compared with mismatch repair-proficient (pMMR) tumors. Additionally, using MSigDB, we compared enrichment scores for hypoxia and DNA repair pathways inside tumor cells, both of which are known to be associated with treatment resistance^18, 30, 31^, across the different pathological response groups; however, no significant differences were observed (Supplementary Fig. 1e). These findings suggest that conventional genomic alterations and treatment resistance pathways in tumor cells alone are insufficient for stratifying patients according to the therapeutic efficacy of CRT.

### CAF and myeloid subsets are associated with therapeutic responses

Given the heterogeneity observed in tumor cell-based analyses, we next investigated the potential contribution of non-tumor cells to treatment resistance. To identify such response- or resistance-associated subpopulations, we employed the Leiden algorithm^32^ to recluster major cell types and identified 10 T cell, 12 myeloid, and 16 stromal cell subpopulations (Fig. 3a, Supplementary Fig. 2). Marker genes for each subcluster were defined through the analysis of differentially expressed genes (DEGs) in each subcluster with respect to all other subclusters within the same major cell type (Supplementary Fig. 2b-d). Among T cell populations, three CD8⁺ T cell subsets (*KLRG1*⁺, *GZMA*⁺, and *BAG3*⁺ T cells) were identified and characterized by elevated expression of cytotoxic markers including *CD8A*, *GZMA*, and *GZMB*. Regulatory T cells (T_regs_) were also identified as a distinct population characterized by high expression of *FOXP3* and *CTLA4*. Among the myeloid populations, *F13A1*⁺ and *IL11*⁺ macrophages (MΦs) expressed high levels of *CD163*, whereas *CD1C*⁺ and *CXCL5*⁺ dendritic cells (DCs) presented elevated expression of *ITGAX*. Within the stromal populations, the CAF markers *FAP* and *PDPN* were upregulated in multiple subclusters, including *PI16*⁺, *SFRP4*⁺, *POSTN*⁺, *MMP11*⁺, *GREM1*⁺, and *CCN1*⁺ CAFs. Additionally, the smooth muscle (SM) marker *MYLK* was upregulated in *LDB*⁺ and *PLN*⁺ stromal subpopulations. Moreover, consistent with previous findings using the Xenium platform^33^, our dataset also revealed the presence of boundary cell populations located at the interfaces between distinct cell types, such as c3_Epi_Mye_boundary, c5_CAF_Mye_boundary, and c0_Epi_CAF_boundary cells (Fig. 3a). To capture potential cell–cell interactions occurring at these interfaces, we retained these boundary cells for downstream analyses.

**Fig. 3.**
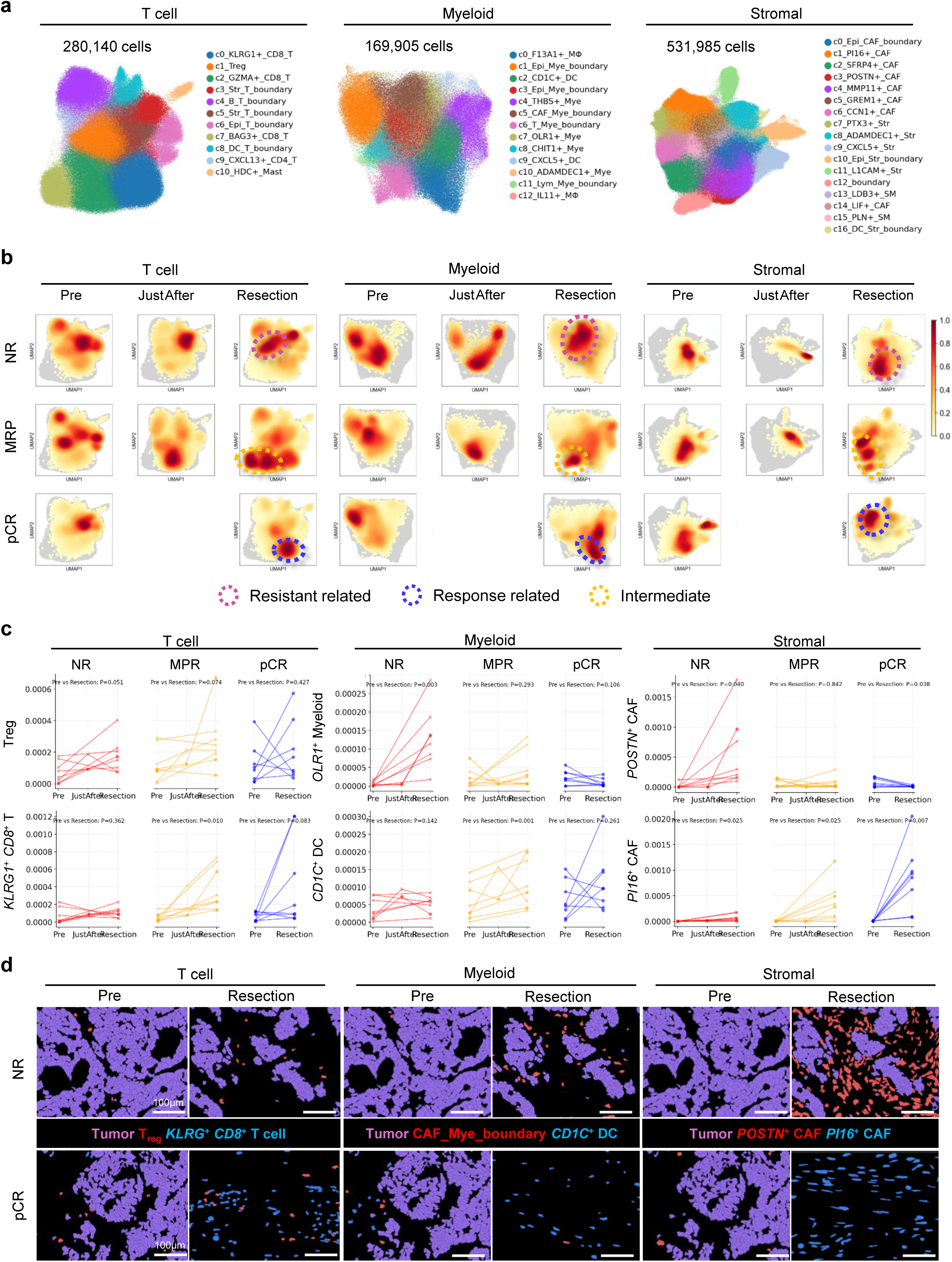
Identification and spatial dynamics of treatment-associated T cells, cancer-associated fibroblasts (CAFs), and myeloid cell subclusters. a, Two-dimensional uniform manifold approximation and projection (UMAP) plots showing subclusters of T cells, myeloid cells, and stromal cells. The subcluster prefixes correspond to the Leiden cluster numbers shown in Supplementary Fig. 2a. b, Density plots in UMAP space for each of the three cell types, stratified by sampling time (Pre, JustAfter, Resection) and treatment response group (NR, MPR, pCR). At each timepoint, regions of high cell density specific to the NR or pCR groups are outlined with red or blue dashed lines, respectively. c, Line plots showing temporal dynamics in cell density across treatment groups for representative subclusters: T cells (regulatory T cells [Tregs], KLRG1⁺ CD8⁺ T cells), myeloid cells (OLR1⁺ myeloid cells, CD1C⁺ dendritic cells), and stromal cells (POSTN⁺ CAFs, PI16⁺ CAFs). Each line represents one patient (NR, red; MPR, yellow; pCR, blue). d, Representative spatial distribution maps of selected response-associated subclusters in NR and pCR patients at the Pre and Resection timepoints. Tumor cells are shown in purple; subclusters enriched in NR patients are highlighted in red, whereas those enriched in pCR patients are shown in blue.

We stratified the UMAP embeddings of T, stromal, and myeloid cells by timepoint and patient clinical response to investigate how CRT shaped immune and stromal cell subpopulations. This analysis revealed distinct subclusters that were specifically enriched posttreatment in the NR, MPR, and pCR groups (Fig. 3b). Within the T cell compartment, the resistant-related subcluster enriched in NR patients was predominantly composed of effector CD8 T cells, whereas the response-related subcluster enriched in pCR patients was mainly composed of T_regs_ (Fig. 3b; Supplementary Fig. 2e). In MPR patients, the intermediate subcluster was primarily characterized by exhausted CD8 T cells. These distributions showed reasonable concordance between treatment responses and T cell phenotypes. We next sought to quantify the density of individual subclusters at the patient level and compare their dynamics across timepoints and pathological response groups. In the NR group, the cell density of *OLR1*⁺ myeloid cells (P = 0.003) and *POSTN*⁺ CAFs (P = 0.040) significantly increased from the Pre to the Resection timepoints, while T_regs_ showed a trend toward increase (P = 0.051) (Fig. 3a–c). In contrast, a significant increase in cell density of *PI16*⁺ CAFs (P = 0.007) was identified in the pCR group, while *KLRG1*⁺ *CD8*⁺ T cells (P = 0.083) and *CD1C*⁺ DCs (P = 0.26) showed non-significant increases. These trends were also observed when cell numbers were expressed as a proportion of total cells or as raw cell counts (Supplementary Fig. 3a, b). At the Resection timepoint, *POSTN*⁺ CAFs (P = 0.036), *MMP11*⁺ CAFs (P = 0.005), c0_Epi_CAF_boundary cells (P = 0.002), *OLR1*⁺ myeloid cells (P = 0.004), c3_Epi_Mye_boundary cells (P = 0.003), and c5_CAF_Mye_boundary cells (P = 0.002) were significantly abundant in the NR group than in the pCR group, whereas *PI16*⁺ CAFs (P = 0.010) were significantly enriched in the pCR group (Supplementary Fig. 3c). Spatial distribution maps of selected subclusters across NR and pCR cases are shown in Fig. 3d, confirming histologically that *POSTN*⁺ CAFs were prominently enriched in NR patients at the post-resection timepoint. Collectively, our findings demonstrate that CRT dynamically alters TME subclusters, revealing distinct CAF and myeloid cell subsets associated with NR (*POSTN*⁺ CAFs, *MMP11*⁺ CAFs, c0_Epi_CAF_boundary cells, *OLR1*⁺ myeloid cells, c3_Epi_Mye_boundary cells, and c5_CAF_Mye_boundary cells) or pCR (*PI16*⁺ CAFs).

### Spatial omics analysis reveals resistance-associated cell populations adjacent to residual tumors

We hypothesized that cell populations contributing to treatment resistance are likely to directly interact with tumor cells. To address this hypothesis, we investigated the dominant cell types located adjacent to residual tumor cells in post-treatment resection samples from all NR patients (n = 8). Specifically, we determined the number of cells from each subcluster located immediately adjacent to the tumor region. The results revealed that *POSTN*⁺ CAFs, *MMP11*⁺ CAFs, c0_Epi_CAF_boundary cells, *OLR1*⁺ myeloid cells, c3_Epi_Mye_boundary cells, and c5_CAF_Mye_boundary cells were the most enriched near tumor cells (Fig. 4a). These subclusters frequently colocalized with each other, suggesting potential cooperative interactions among these cell types at the tumor–stroma interface (Fig. 4b, c). We then defined NR-associated CAFs (NR CAFs) as *POSTN*⁺ CAFs, *MMP11*⁺ CAFs, and c0_Epi_CAF_boundary cells, and NR-associated myeloid cells (NR myeloid cells) as *OLR1*⁺ myeloid cells, c3_Epi_Mye_boundary cells, and c5_CAF_Mye_boundary cells. The abundance of these NR CAF and NR myeloid cell populations at the resection timepoint was significantly higher in NR cases than in both pCR and MPR patients (Supplementary Fig. 4a). Moreover, in NR patients, these populations were significantly enriched in regions adjacent to tumor cells (Supplementary Fig. 4b). To elucidate cell–cell communication among these NR-associated cells adjacent to residual tumor cells, we applied CrossTalkeR to infer ligand–receptor interactions (Supplementary Fig. 4c). This analysis, performed within 30 μm of tumor cells using all ligand–receptor pairs from the LIANA consensus database, identified multiple significant interactions across these cellular compartments. Among the most highly ranked of these, POSTN expressed by NR CAFs was predicted to engage ITGAV/ITGB5 expressed by NR CAFs, NR myeloid cells, and tumor cells, and NR myeloid cells and NR CAFs were predicted to signal via PLAU, targeting PLAUR, IGF2R, LRP1, and ITGB5 in neighboring cell populations (Supplementary Fig. 4d). Together, these observations support a tumor-promoting and immunosuppressive residual tumor microenvironment.

**Fig. 4.**
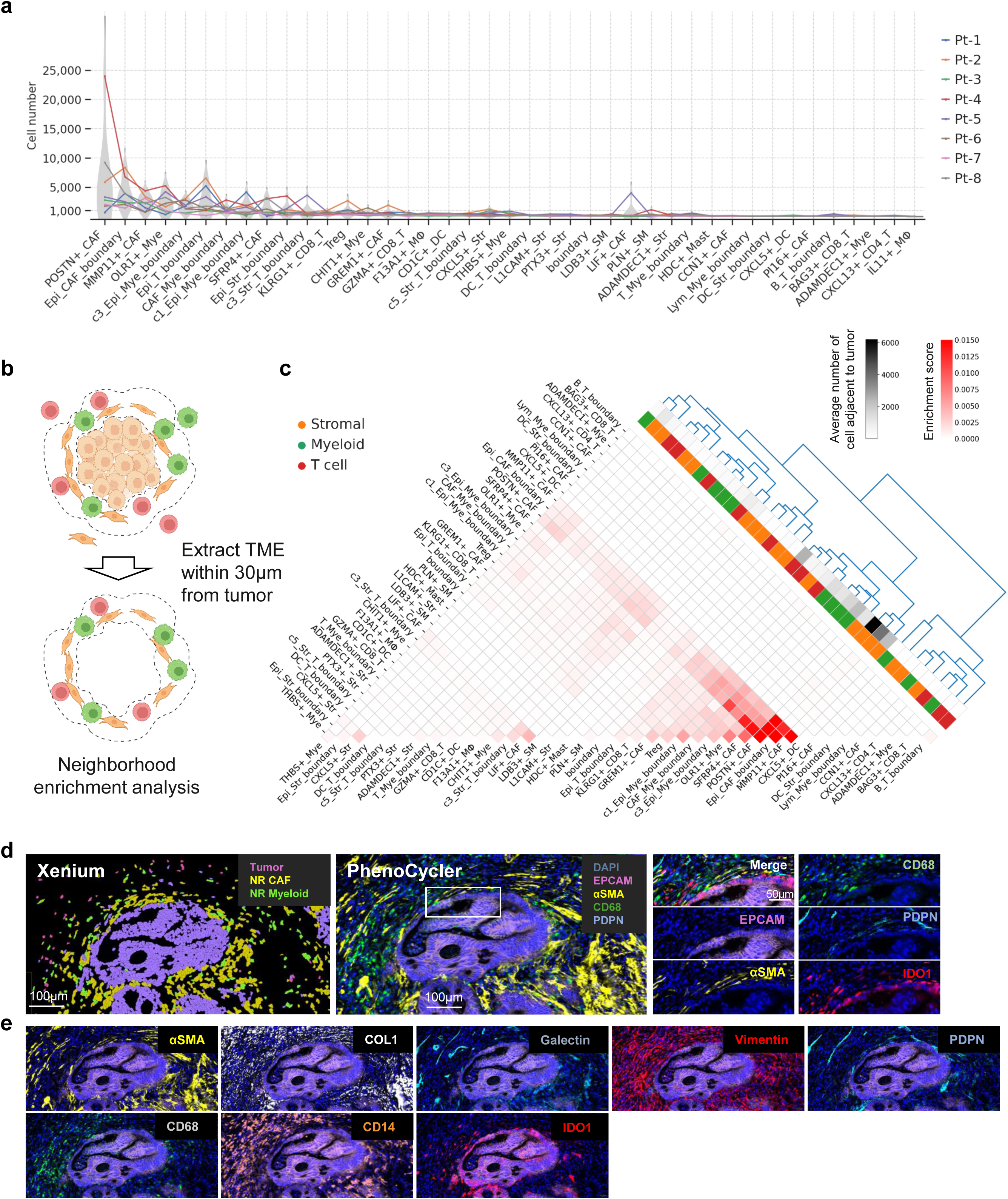
Treatment-resistant CAFs and myeloid cells are abundant in the tumor-adjacent microenvironment. a, Line plots showing the number of cells adjacent to tumor cells across various subclusters. Each line color represents one of the eight NR patients. Subclusters on the x-axis are ordered by descending mean adjacency across the NR cohort. b, Schematic illustration of neighborhood enrichment analysis. Tumor microenvironment (TME) regions within 30 μm of tumor cells were extracted from Xenium data, and spatial neighborhood patterns were analyzed using the SpatialKNifeY (SKNY) algorithm^52^. c, Upper triangular clustered heatmap representing the spatial proximity among subclusters within 30 μm of tumor cells. Hierarchical clustering was performed using Ward’s algorithm with the Euclidean distance. The heatmap color intensity (red scale) indicates the average proportion of adjacent cell pairs across the eight NR patients. The categorical annotation bar represents major cell types, whereas the grayscale bar indicates the average number of tumor-adjacent cells per subcluster in the NR group. The blue dendrogram shows the distance based on the similarity of adjacency counts among different cell types. d, Spatial localization of NR-enriched CAFs and myeloid cells identified by Xenium (left), corresponding protein expression patterns determined by PhenoCycler on the same tissue sections (middle), and magnified views (right). In the Xenium panel, purple indicates tumor cells, yellow indicates NR CAFs, and green indicates NR myeloid cells. PhenoCycler images show the expression of DAPI, EPCAM, αSMA, CD68, PDPN, and IDO1. e, Representative protein expression visualized with the PhenoCycler platform. The upper row shows CAF-related markers (αSMA, collagen I, galectin, vimentin, and PDPN), and the lower row shows macrophage-related markers (CD68, CD14, and IDO1).

To further decipher the phenotypes of these NR-associated cells, we compared the spatial transcriptomic localization obtained on the Xenium platform with the protein expression detected in the same section using the PhenoCycler platform (Fig. 4d, e). NR CAFs were found to strongly express αSMA, collagen I (COL1), galectin, vimentin, and podoplanin (PDPN), which is consistent with a myofibroblastic CAF (myCAF)-like phenotype and has been associated with extracellular matrix remodeling and tumor progression^34, 35^. NR myeloid cells were shown to express CD68, CD14, and the immunosuppressive marker IDO1, which are indicative of an immunosuppressive tumor-associated macrophage phenotype. These findings demonstrate that both NR CAFs and myeloid cells exhibit phenotypes associated with treatment resistance. Our analysis revealed that, among the diverse TME components examined, specific functional subsets of CAFs and myeloid cells constitute the key determinants of CRT resistance.

Tumor and normal epithelial regions were identified by a pathologist based on H&E staining, and we confirmed that NR CAFs were specifically localized around tumor cells and largely absent from normal epithelial tissues (Fig. 5a). In contrast, normal fibroblasts were localized exclusively within the normal intestinal epithelium (Fig. 5b; Supplementary Fig. 5). pCR-associated CAFs (pCR CAFs), defined as *PI16*⁺ CAFs, were predominantly localized in tumor bed areas where tumor cells had regressed (Fig. 5c). We examined the expression of CAF marker proteins to characterize the distinct features of these CAF subsets using the PhenoCycler system. NR CAFs showed strong αSMA and PDPN expression, whereas collagen I signals were relatively weak (Fig. 5d). In contrast, pCR CAFs displayed high collagen I expression and intermediate galectin-3 expression. Moreover, NR CAFs and normal fibroblasts appeared morphologically rounder than pCR CAFs (Fig. 5d). The distinct spatial localization and phenotypes of NR CAFs relative to normal fibroblasts and pCR CAFs suggest that NR CAFs constitute a unique population, potentially enabling selective targeting of NR CAFs while minimizing off-target effects on normal tissue.

**Fig. 5.**
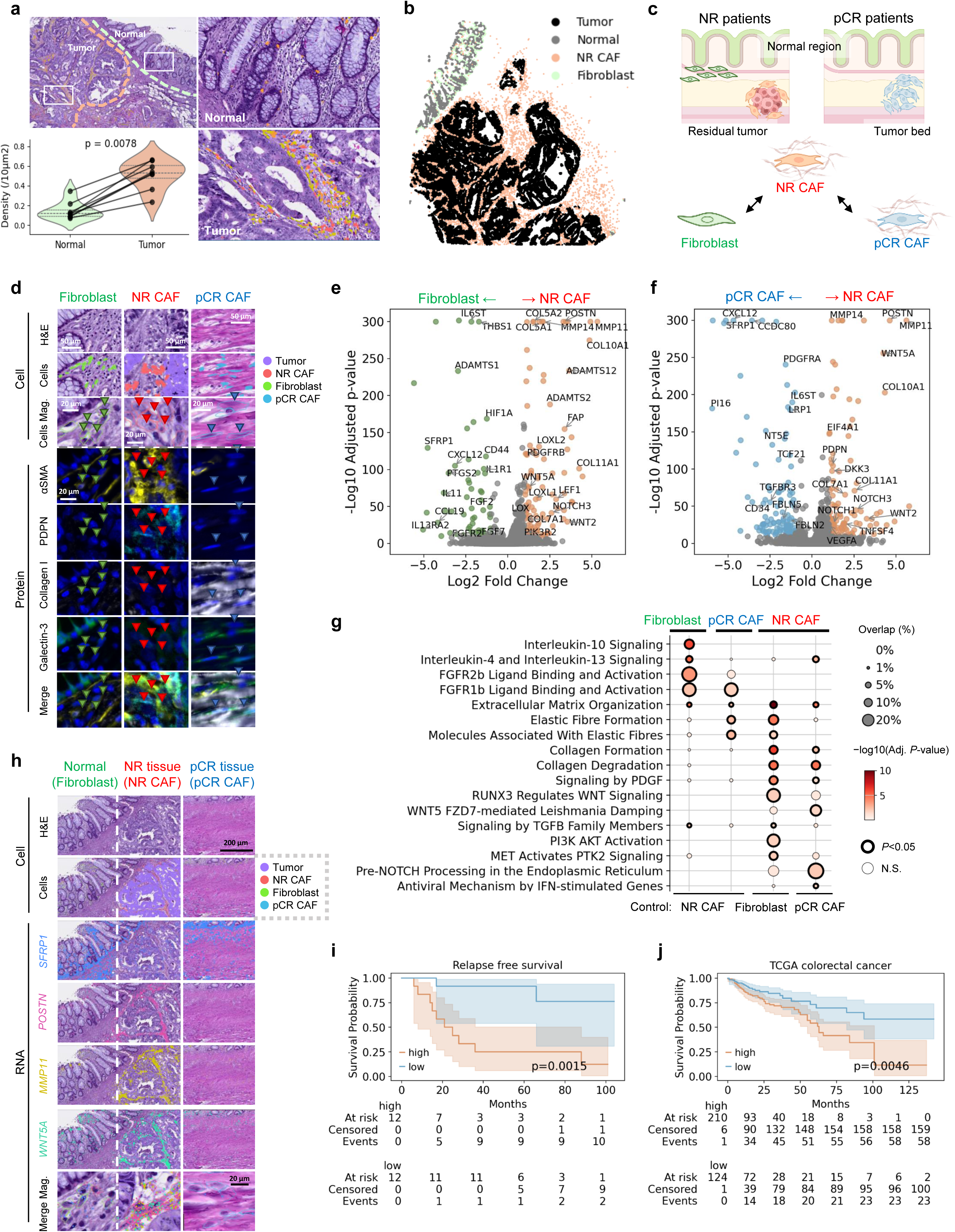
Molecular characteristics and targetable pathways of NR CAFs. a, Spatial visualization of nonresponder cancer-associated fibroblasts (NR CAFs) within both tumor and normal epithelial regions. The top-left panel shows a hematoxylin and eosin (H&E)-stained image merged with the spatial distribution of NR CAFs (red, orange, and yellow). The right panels show magnified views of the normal epithelial region (top) and the tumor region (bottom). The bottom-left violin plot displays the density of NR CAFs within 30 μm of tumor and normal epithelial cells (see Methods). Each line represents one of the eight NR patients. The P value was calculated using the paired t test. b, Spatial distribution of tumor cells (black), normal epithelial cells (gray), NR CAFs (red), and fibroblasts (blue) in patients Pt-2 and Pt-7. The data for the remaining six NR patients are shown in Supplementary Fig. 5. c, Summary schematic of the comparative analyses. To characterize the gene expression profiles and morphological features of NR CAFs, fibroblasts adjacent to the normal epithelia of NR group patients (n = 8) and pCR-dominant CAFs from pCR group patients (n = 8; see Fig. 3c) were used as controls. Comparative analyses included morphology (H&E staining), protein expression (PhenoCycler), gene expression and spatial distribution (Xenium), and survival outcomes (relapse-free survival and overall survival). d, Comparison among fibroblasts, NR CAFs, and pCR CAFs across multiple modalities: row 1, H&E staining; rows 2–3, Xenium-based spatial distribution; rows 4–8, protein expression by PhenoCycler. The green, red, and blue arrowheads indicate fibroblasts, NR CAFs, and pCR CAFs, respectively. e, f, Differential gene expression (DEG) analysis comparing fibroblasts (control) and NR CAFs (e) and pCR CAFs (control) and NR CAFs (f). Volcano plots show genes upregulated in NR CAFs (red) and controls (fibroblasts, green; pCR CAFs, blue). The DEG thresholds were set to P < 1×10⁻¹⁰ and |log₂ fold change| > 1. g, Pathway enrichment analysis using the Reactome pathway database. The color scale indicates -log₁₀(FDR). The size of the dots indicates the overlap between the upregulated genes and each pathway. h, Comparison of spatial localization across fibroblasts, NR CAFs, and pCR CAFs. Row 1: H&E staining; row 2: spatial distribution and merged images from Xenium; rows 3–5: Xenium-based spatial localization of SFRP1, POSTN, and MMP11 with H&E overlay. Fibroblast and pCR CAF regions were taken from adjacent areas within the same tissue section. i, Kaplan–Meier curve of relapse-free survival in patients grouped by total POSTN expression per tissue section (high versus low). j, Kaplan–Meier curve of overall survival in the TCGA CRC cohort stratified by POSTN expression. The red and blue lines indicate the high-and low-expression groups, respectively. P values were calculated using the log-rank test.

### Transcriptome analysis uncovers therapeutic targets in NR CAFs

Having shown that NR CAFs are specifically localized adjacent to residual tumor cells, indicating their promise as selective therapeutic targets, we performed DEG analysis using Xenium data to identify molecular targets underlying their pathogenic role. Compared with normal fibroblasts, NR CAFs showed significant upregulation of matrix remodeling genes, including *POSTN*, *COL10A1*, *COL5A1*, *MMP11*, *MMP14*, and *SULF1* (Fig. 5e, g). Moreover, compared with pCR CAFs, NR CAFs exhibited upregulation of *WNT5A* and *COL4A1* and downregulation of *CXCL12*, *SFRP1*, and *PI16* (Fig. 5e, f). Reactome pathway analysis revealed distinct gene expression profiles among normal fibroblasts, pCR CAFs, and NR CAFs. Compared to NR CAFs, normal fibroblasts and pCR CAFs exhibited enrichment of tumor-suppressive pathways, including interleukin-10 signaling, FGFR1b/FGFR2b signaling, and elastic fiber formation (Fig. 5g). In contrast, NR CAFs exhibited enrichment of tumor-promoting pathways, such as collagen formation, collagen degradation, WNT signaling, TGFB signaling, PI3K/Akt signaling, Notch signaling, and IFN signaling.

We next sought to validate that the pathways upregulated in NR CAFs were localized around residual tumor cells through pathology image analysis. *SFRP1*, a fibroblast-specific gene, was found in normal epithelial cells, whereas *POSTN*, *WNT5A*, and *MMP11*—genes upregulated in NR CAFs—were specifically localized around residual tumor cells (Fig. 5h). Given the tumor-specific spatial localization of POSTN, we sought to explore whether this expression was associated with treatment resistance. In our study cohort, Kaplan–Meier survival analysis based on total *POSTN* expression per tissue section demonstrated that high expression was significantly associated with reduced relapse-free survival (RFS; P = 0.0015) (Fig. 5i). Validation in an independent The Cancer Genome Atlas (TCGA) CRC cohort confirmed the association between elevated *POSTN* expression and shorter overall survival (OS; P = 0.0046) (Fig. 5j). These results suggest that *POSTN*-expressing NR CAFs contribute to resistance to CRT and promote tumor recurrence.

### Trajectory analysis suggests therapy-induced bifurcation into NR and pCR CAF states

NR and pCR CAFs were predominantly observed at the Resection timepoint, suggesting that these represent treatment-associated CAF states. To explore their potential origin, we performed trajectory inference using the STORIES algorithm on CAF subclusters from both Pre and Resection samples. This analysis revealed a bifurcating trajectory in which Pre CAFs diverged into NR and pCR branches after CRT, with CAFs from both NR and pCR Pre samples positioned at the origin of the trajectory (Fig. 6a,b; Supplementary Fig. 6a). In addition, using the same approach as previously reported, we estimated the fate probabilities of Pre CAFs toward NR and pCR CAF states. The inferred probabilities were comparable between NR and pCR patients and showed no significant difference (Fig. 6c,d). Taken together, these results suggest that NR and pCR CAFs originate from a common Pre CAF population and that commitment toward either state is not predetermined at the pretreatment stage.

**Fig. 6.**
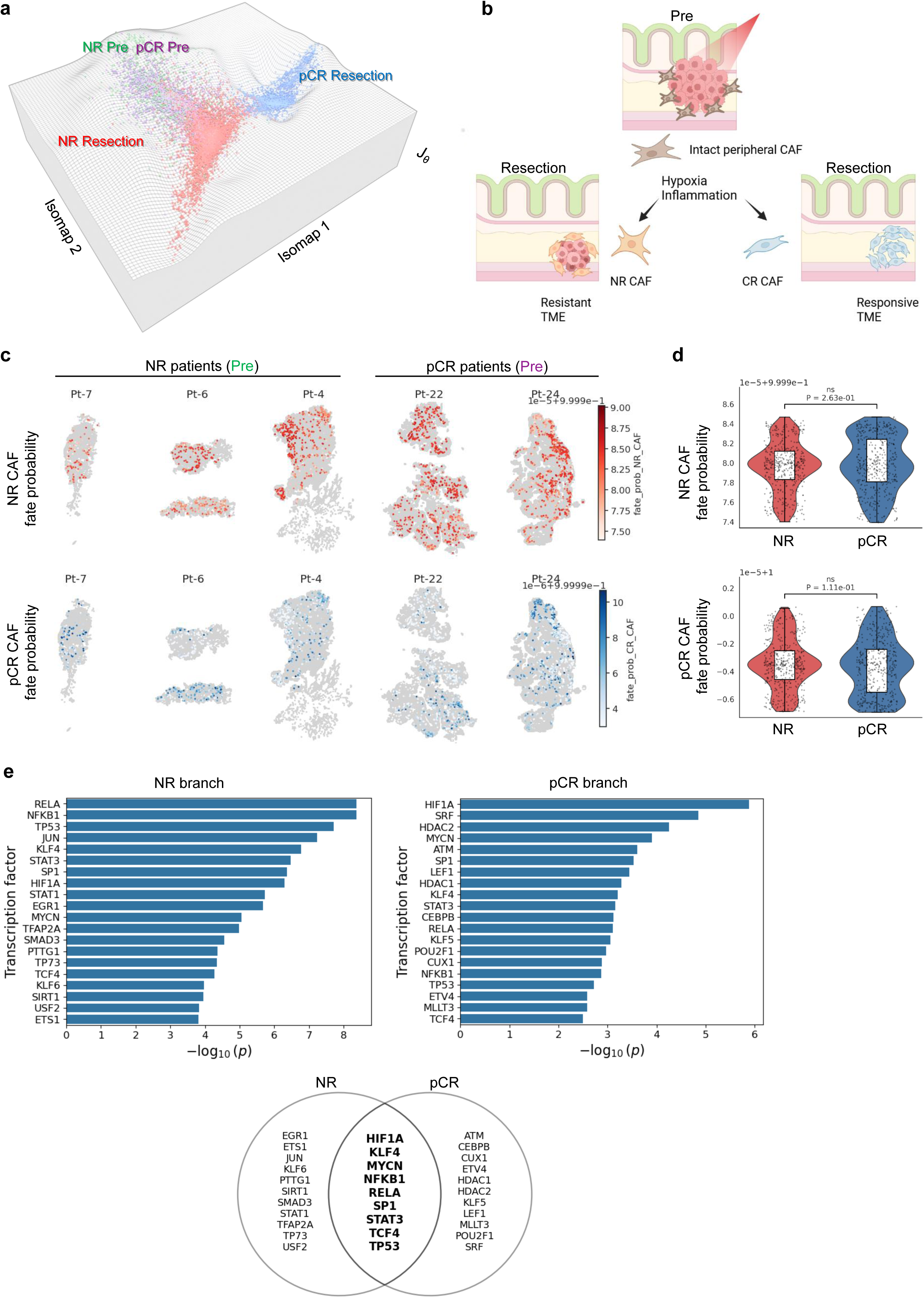
Trajectory inference reveals bifurcation of pretreatment CAFs into NR and pCR branches after CRT. **a**, Trajectory inference of CAFs using the STORIES algorithm. CAFs from Pre and Resection samples were embedded using Isomap, revealing a bifurcating trajectory originating from Pre CAFs and diverging into NR and pCR branches after CRT. Cells are colored according to clinical response and sampling timepoint (NR Pre, pCR Pre, NR Resection, and pCR Resection). **b**, Schematic model illustrating the inferred differentiation trajectory of CAFs. Intact peripheral CAFs present at the Pre timepoint bifurcate into two transcriptional states following CRT. **c**, Visualization of inferred fate probabilities for individual Pre CAFs toward NR CAF or pCR CAF states across representative patients. Cells are colored according to the predicted probability of differentiation toward NR CAF (top) or pCR CAF (bottom) states. **d**, Comparison of predicted fate probabilities of Pre CAFs toward NR CAF or pCR CAF states between NR and pCR patients. Boxplots represent the median and interquartile range; P values were calculated using the Mann–Whitney U test. **e**, Transcription factor enrichment analysis along the NR and pCR branches using TRRUST. Bar plots show enriched transcription factors ranked by –log10(P value). The Venn diagram summarizes transcription factors shared between the two branches and those uniquely enriched in each branch.

To investigate how this common progenitor population diverges into NR CAFs and pCR CAFs, we next examined genes whose expression changed along pseudotime in the NR and pCR branches. In the NR branch, expression of genes associated with extracellular matrix remodeling and cytoskeletal regulation, including POSTN, RHOC, and RAC1, increased along pseudotime (Supplementary Fig. 6b). In contrast, in the pCR branch, genes such as WNT5A and MMP11 showed a decreasing trend (Supplementary Fig. 6c). To identify transcriptional regulators potentially driving these transcriptional changes, we performed transcription factor enrichment analysis using TRRUST. Several transcription factors were commonly enriched in both branches, including HIF1A, KLF4, MYCN, SP1, STAT3, TP53, RELA, and NFKB1, suggesting shared regulatory programs related to stress responses and inflammatory signaling that likely represent a common transcriptional response of CAFs to CRT (Fig. 6e). In addition, the NR branch showed enrichment of transcription factors such as JUN, SMAD3, STAT1, EGR1, and ETS1, which are associated with inflammatory signaling, TGF-β-related transcriptional regulation, and cellular activation. In contrast, the pCR branch was characterized by enrichment of transcription factors including LEF1, CEBPB, SRF, HDAC1, and HDAC2, suggesting involvement of WNT-related transcriptional programs and chromatin regulatory mechanisms. Collectively, these findings suggest that differentiation toward the NR CAF state may be driven, at least in part, by TGF-β-related transcriptional programs.

### NR CAFs promote pro-tumor recurrence states

Because NR CAFs are more abundant around residual tumor cells and express both secreted paracrine factors (such as WNT5A and TNFSF4) and structural proteins (such as COL10A1 and COL5A1), we reasoned that neighboring tumor cells would exhibit distinct molecular functions. To investigate the potential influence of NR CAFs on the tumor cell state, we categorized tumor cells as “neighbor” or “nonneighbor” depending on whether they made direct contact with NR CAFs (Fig. 7a, b; Supplementary Fig. 7a). DEG analysis revealed that CAF-neighbor tumor cells exhibited upregulated expression of genes associated with cell adhesion, remodeling, and inflammatory signaling, including ITGB4, ITGB6, BMP4, NFKB2, TNFSF6B, TNFSF11B, PDGFB, and LAMC2 (Fig. 7c, d). Gene set enrichment analysis via MSigDB revealed significant activation of EMT and TNF-α signaling via NF-κB in all patients (Fig. 7e). These pathways are strongly implicated in CRC progression, recurrence, and resistance to therapy^36, 37^. We also observed activation of several EMT-associated pathways—including Notch signaling, TGF-β signaling, IL-6/JAK/STAT3 signaling, Wnt/β-catenin signaling, noncanonical Wnt signaling, IL-2/STAT5 signaling, hypoxia, and mTORC1 signaling—in neighboring tumor cells. The activation patterns of these pathways exhibited substantial interpatient heterogeneity compared with that of EMT (Fig. 7e; Supplementary Fig. 7b). These findings demonstrate that NR CAFs drive EMT in adjacent tumor cells through patient-specific multipathway mechanisms.

**Fig. 7.**
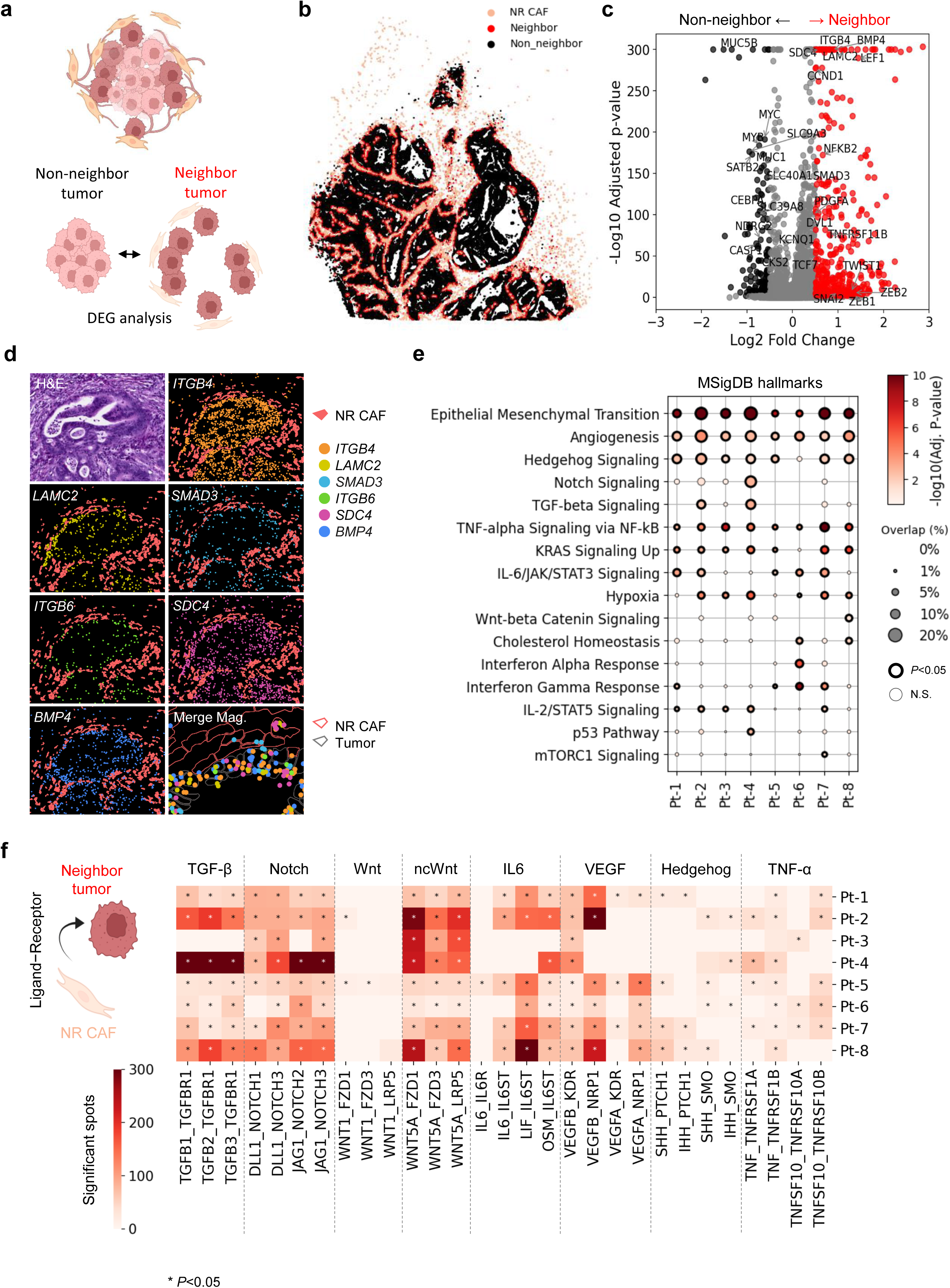
Molecular function of tumor cells adjacent to NR CAFs. a, Schematic diagram of the workflow used to classify tumor cells as either neighboring or nonneighboring NR CAFs (see Methods). Differentially expressed gene (DEG) analysis was conducted between these groups to identify the molecular functions associated with tumor cells near NR CAFs. b, Identification of tumor cells neighboring NR CAFs using the spatial neighborhood graph algorithm in Squidpy (see Methods). The spatial locations of neighboring (red) and nonneighboring (gray) tumor cells in patient Pt-7 are shown. The data for the other seven NR patients are presented in Supplementary Fig. 7a. c, Volcano plot showing the results of DEG analysis between neighboring and nonneighboring tumor cells. Genes upregulated in neighboring tumor cells are shown in red, whereas those upregulated in nonneighboring tumor cells are shown in black. Thresholds: P < 0.01 and |log₂ fold change| > 0.5. d, Representative hematoxylin and eosin (H&E)-stained image containing tumor-stroma boundaries and spatial localization of selected genes detected by Xenium: ITGB4 (orange), LAMC2 (yellow), SMAD3 (cyan), ITGB6 (green), SDC4 (magenta), and BMP4 (blue). High-magnification merged images with cell segmentation masks (gray) are also shown. e, Pathway enrichment analysis of genes upregulated in neighboring tumor cells using the Molecular Signatures Database (MSigDB). The color scale (white to red) indicates -log₁₀(FDR). The size of the dots indicates the overlap between the upregulated genes and each pathway. f, Heatmap showing ligand–receptor interactions between NR CAFs and neighboring tumor cells. * indicates P < 0.05.

We next aimed to identify the ligand–receptor interactions mediating CAF–tumor cell communication associated with these activated signaling pathways in the TME. By performing ligand–receptor interaction analyses among NR CAFs, normal fibroblasts, pCR CAFs, neighbor tumor cells, and nonneighbor tumor cells, we identified significant interactions between NR CAFs and neighbor tumor cells (Supplementary Fig. 7c). Although interactions involving angiogenesis and Hedgehog signaling were not significant, we identified notable ligand–receptor pairs linked to Wnt signaling (e.g., WNT5A–FZD1 and WNT5A–LRP6), TGF-β signaling (e.g., TGFB1/3–TGFBR1/2), TNF-α signaling via NF-κB (e.g., TNFSF10–TNFRSF10A/B and TNF–TNFRSF1A), Notch signaling (e.g., DLL1/JAG1–NOTCH1/2), and IL-6/JAK/STAT3 signaling (IL6–IL6ST). Collectively, these results suggest that the promotion of tumor EMT by NR CAFs is a dominant phenomenon across all patients, whereas the upstream signaling pathways and specific ligand–receptor pairs involved demonstrate considerable interpatient variability (Fig. 7f). In summary, single-cell spatial analysis revealed common myeloid and CAF subpopulations responsible for establishing a CRT-resistant microenvironment. In particular, NR CAFs emerged as central mediators that promote recurrence-associated programs in residual tumor cells through multiple signaling pathways, highlighting their potential as rational therapeutic targets (Supplementary Fig. 8).

## Discussion

In this study, we utilized single-cell spatial transcriptomics to comprehensively characterize the tumor microenvironment (TME) after chemoradiotherapy (CRT) in patients with locally advanced rectal cancer (LARC). By focusing on previously reported mediators of radioresistance across cancer, immune, and stromal compartments, we found that CAF-mediated mechanisms are dominant within patient tissues and are consistently activated in treatment-resistant cases. In addition to these mechanistic insights, the paired pre- and post-CRT Xenium and PhenoCycler profiles, together with matched Visium HD datasets generated on adjacent serial sections, provide a clinically annotated multimodal resource for interrogating CRT-induced spatial remodeling.

From a Xenium dataset comprising approximately 2.9 million cells, we identified distinct cell clusters associated with therapeutic response and resistance. Notably, tumor cell clusters displayed substantial interpatient variability, and we did not observe tumor cell clusters that were consistently associated with treatment responses across patients. These findings highlighted the importance of investigating resistance mechanisms conserved among patients. We therefore focused on resistance-associated non-tumor cell populations within the TME. This analysis identified regulatory T cells (Tregs), *OLR1***^+^**myeloid cells, and *POSTN***^+^** CAF subclusters associated with non-response (NR). Among these, NR-associated CAF clusters were particularly notable, as they were consistently enriched and accumulated adjacent to residual tumor cells across nearly all treatment-resistant cases.

Using the Xenium dataset, we next characterized the spatial, temporal, and functional properties of NR-associated CAFs (NR CAFs). NR CAFs were rarely detected before treatment and were largely absent from normal intestinal mucosa, but they newly emerged after CRT and were enriched around residual tumor cells, indicating that NR CAFs represent a therapy-induced cell population. Trajectory inference further suggested that pretreatment tumor-peripheral CAFs bifurcate toward both NR CAF and pCR-associated CAF (pCR CAF) states after CRT (Fig. 6a, b; Supplementary Fig. 6), supporting CAF remodeling under radiotherapy-associated stress (e.g., hypoxia and inflammatory stress) as a plausible origin of these treatment-associated CAF states. Consistently, transcription factor enrichment analyses suggested broadly shared stress-response regulators across both branches, including HIF1A, NF-κB components, and TP53 (Fig. 6e), whereas clear NR CAF-specific upstream signals were limited in this analysis. Although defining NR CAF-specific cues and vulnerabilities remains an important future direction, the posttreatment-specific emergence of NR CAFs and their scarcity in normal tissues support their potential as highly specific therapeutic targets.

To define the molecular features of NR CAFs, we compared NR CAFs with normal fibroblasts and pCR CAFs. In pCR CAFs, we observed features consistent with previously reported tumor-suppressive CAF phenotypes, including PI16 and type I collagen expression^38, 39^. In contrast, NR CAFs exhibited markedly reduced collagen protein levels, despite increased collagen gene expression relative to normal fibroblasts, suggesting enhanced matrix metalloproteinase (MMP)-dependent extracellular matrix remodeling. We further identified MMP11, WNT5A, and POSTN as characteristic markers of NR CAFs. Although elevated POSTN expression has been associated with poor prognosis across multiple cancer types, prior studies have predominantly focused on POSTN functions in cancer cells, and its role in CAFs has remained poorly defined^40^. Our data highlight that POSTN is highly expressed specifically in NR CAFs in a subset of LARC patients after CRT, emphasizing the clinical relevance of *POSTN***^+^** CAFs in posttreatment resistant tumor ecosystems.

Single-cell spatial analyses further demonstrated that, across all treatment-resistant cases, residual tumor cells adjacent to NR CAFs displayed induction of epithelial–mesenchymal transition (EMT). Multiple signaling pathways were implicated in NR CAF–tumor interactions, including Notch, TGF-β, IL-6/JAK/STAT3, Wnt/β-catenin, IL-2/STAT5, hypoxia, and mTORC1 signaling^41, 42, 43^; however, their activation patterns varied among patients. Notably, Pt-3—the only patient in the NR group who did not experience recurrence—showed the weakest activation of these signaling interactions. In contrast, EMT and angiogenesis were consistently enriched across treatment-resistant cases, indicating that NR CAFs and EMT constitute central and conserved features of CRT resistance despite interpatient variability in upstream signaling.

To date, numerous therapeutic strategies have attempted to overcome resistance by targeting cancer cell–intrinsic pathways such as WNT signaling, MMPs, and integrins, yet clinical efficacy has been limited^44, 45, 46, 47^. This limited efficacy likely reflects the complex and redundant signaling networks supporting resistance. Our study suggests that NR CAFs may emerge as a central regulatory node in resistant tumors and that targeting POSTN+ CAFs may provide a more comprehensive strategy by simultaneously attenuating multiple downstream pro-resistance pathways. POSTN-targeted therapies have been actively investigated in inflammatory diseases, and multiple POSTN-targeting antibodies have already been evaluated in preclinical settings^48, 49, 50^, supporting the feasibility of translational development.

This study has several limitations. First, the sample size was limited, precluding a comprehensive evaluation of how genetic backgrounds—such as mismatch repair status or driver mutations—shape treatment responses. Second, our analysis was restricted to a predefined panel of approximately 5,000 genes. Future studies leveraging transcriptome-wide platforms such as Visium HD and integrative analyses with complementary datasets may provide additional insights. We anticipate that this clinically annotated multimodal atlas will serve as a reference for studying radiotherapy-driven TME reprogramming and for developing and benchmarking hypotheses in future mechanistic and translational studies. Given the widespread use of neoadjuvant CRT in LARC and the active development of CRT-based combination regimens, this atlas provides an important foundation for these efforts by enabling comprehensive interrogation of post-CRT tissue remodeling in situ^10, 11, 12, 13, 14, 15^.

In conclusion, our single-cell spatial transcriptomics analysis provides new insights into the molecular and cellular basis of CRT resistance in LARC and identifies therapy-induced *POSTN***^+^** NR CAFs as a central component of resistant tumor microenvironments. Further validation and translational studies are warranted to assess the clinical potential of targeting *POSTN***^+^** CAFs in combination with current treatment modalities.

## Methods

### Sample collection

Samples from tumor tissues and adjacent normal tissues were collected from 24 patients who were diagnosed with LARC and underwent surgical resection at the National Cancer Center Hospital East. For each patient, samples were obtained at two standard timepoints: pretreatment (Pre) and at the time of surgical resection (Resection) (Fig. 1a). Additionally, samples within 2 weeks after completion of CRT (JustAfter) were obtained from 4 patients, as immediate post-treatment biopsy is not part of standard clinical protocol. Pre and JustAfter samples were collected using biopsy forceps, while Resection samples were obtained through surgical procedures. Informed consent was obtained from all participants, and the study was approved by the institutional review board (IRB protocol number: 2018-101).

### In situ gene expression analysis with the Xenium platform

In situ spatial transcriptomic analysis was performed using the Xenium platform (10x Genomics) following the manufacturer’s protocol and optimized for formalin-fixed paraffin-embedded (FFPE) tissue. In brief, FFPE sections (5 μm thick) were mounted on Xenium-compatible slides; after deparaffinization, the sections were subjected to antigen retrieval and protease treatment. Gene expression profiling was carried out using the Human Gene Expression Panel (5,001 genes). To ensure comparability across samples profiled with the default panel alone or with an additional add-on panel, analyses were restricted to the 5,001 genes shared between the panels. Following signal amplification and imaging, the raw spatial transcriptomic data were processed using the Xenium Analyzer pipeline. Single-cell segmentation and transcript count quantification were performed to generate spatially resolved gene expression matrices.

### Spatial transcriptomics with Visium HD

Spatial transcriptomic profiling was additionally performed on adjacent serial sections using the Visium HD Spatial Gene Expression assay (10x Genomics) following the manufacturer’s protocol for FFPE samples. In brief, FFPE sections were deparaffinized, stained and imaged to record tissue morphology, and then processed for probe hybridization and library preparation to generate spatially barcoded gene expression libraries. Sequencing libraries were sequenced on an Illumina platform. Raw sequencing data were processed using Space Ranger (10x Genomics) to generate spatial gene expression count matrices aligned to the GRCh38 reference genome. These Visium HD datasets were generated as a complementary resource and were not included in the primary analyses presented in this study.

### Multiplexed immunostaining with PhenoCycler

Multiplexed immunofluorescence staining and imaging were performed using the PhenoCycler platform (Akoya Biosciences). Tissue sections were stained with a panel of validated antibodies conjugated to oligonucleotide barcodes (Supplementary Table 1). After multiple rounds of hybridization and imaging, signal deconvolution was performed to generate multiplexed protein images. Because this assay was performed in combination with Xenium, image quality was substantially reduced compared with conventional PhenoCycler imaging. Therefore, the PhenoCycler data were used primarily for qualitative comparison of spatial protein localization with Xenium results, whereas robust cell segmentation and quantitative signal measurement were not feasible in this dataset.

### Hematoxylin and eosin (H&E) staining

Consecutive tissue sections were stained with hematoxylin and eosin (H&E) following standard histopathological protocols. Images were acquired using a high-resolution whole-slide scanner (e.g., Hamamatsu NanoZoomer). Then, a certified pathologist manually delineated the tumor regions and normal epithelium on the images of each slide. These annotations were used to guide downstream spatial transcriptomic and proteomic analyses.

### Whole-exome sequencing (WES)

Genomic DNA was extracted from tumor and matched normal tissue samples using a QIAamp DNA Mini Kit (Qiagen). Whole-exome libraries were prepared using a SureSelect Human All Exon V7 Kit (Agilent) and sequenced on an Illumina NovaSeq platform with 150-bp paired-end reads; the resulting average coverages were >100× for tumor samples and >50× for normal samples. Quality control metrics were assessed with FastQC and MultiQC.

### Variant calling using whole-exome sequencing data

The sequencing reads were aligned to the human reference genome (GRCh38) with BWA-MEM. PCR duplicates were marked using Picard, and base quality recalibration and realignment were performed using GATK. Somatic variants were identified with Mutect2 (GATK v4) by comparing the reads from the tumor and matched normal samples. Variants were filtered with the FilterMutectCalls module of GATK, and variants were annotated with their functional consequences with ANNOVAR and VEP. Variants in cancer-related genes (e.g., *KRAS*, *PIK3CA*, and *TP53*) were manually reviewed and employed in downstream correlation analyses with spatial transcriptomic data.

### Preprocessing of Xenium expression data

Gene expression data from 52 Xenium samples (n = 2,829,869 cells) were loaded as individual AnnData objects and concatenated using Scanpy (v1.9.8). Sample identifiers were retained in the metadata during integration. Normalization was performed using sc.pp.normalize_total, scaling total transcript counts per cell to a common library size (default target sum), followed by logarithmic transformation (sc.pp.log1p). Highly variable genes (HVGs) were identified using sc.pp.highly_variable_genes with n_top_genes = 2000, and principal component analysis (PCA) was performed on the HVGs using sc.tl.pca. A k-nearest neighbor (KNN) graph was constructed based on the PCA embedding using sc.pp.neighbors (default parameters). Uniform Manifold Approximation and Projection (UMAP) was computed for visualization (sc.tl.umap). Clustering was performed using the Leiden algorithm (sc.tl.leiden, default resolution). Quality control metrics were calculated using sc.pp.calculate_qc_metrics. Clusters were ranked according to the median number of detected genes per cell (n_genes_by_counts), and clusters were categorized into high-, middle-, and low-quality groups. Cells belonging to low- and middle-quality clusters were excluded. Additionally, cells expressing fewer than 100 genes or with fewer than 100 total counts were removed from downstream analyses.

### Annotation of major cell clusters

The filtered dataset was reprocessed and reclustered using rapids-singlecell (Scanpy-compatible GPU implementation). Specifically, total counts were normalized (normalize_total), log-transformed (log1p), and highly variable genes were identified (n_top_genes = 2000). PCA was recomputed based on the highly variable genes. A new KNN graph was constructed from the PCA embedding (pp.neighbors, random_state = 0), and clustering was performed using the Leiden algorithm (tl.leiden, random_state = 0). Major cell types—including epithelial cells, stromal cells, T cells, myeloid cells, and B cells—were manually annotated based on canonical marker gene expression. Clusters containing mixed major cell identities were excluded from further analyses.

Tumor and normal epithelial regions were annotated on H&E-stained tissue sections by a board-certified pathologist. Based on these annotations, corresponding cell IDs were extracted using Xenium Explorer, enabling separation of tumor epithelial and normal epithelial cell populations.

### Subcluster cell annotation

Stromal, T cell, and myeloid populations were subset from the high-quality integrated dataset and reclustered independently using Scanpy. For each subset, highly variable genes were identified (sc.pp.highly_variable_genes, n_top_genes = 2000), followed by principal component analysis (sc.tl.pca). A KNN graph was constructed based on the PCA embedding using default parameters (sc.pp.neighbors), and dimensionality reduction for visualization was performed using UMAP (sc.tl.umap). Clustering was performed using the Leiden algorithm (sc.tl.leiden, default resolution). Differentially expressed genes (DEGs) for each subcluster were identified using the Wilcoxon rank-sum test (sc.tl.rank_genes_groups, method = "wilcoxon", each cluster compared against all others). Genes with adjusted *P* values below 1×10⁻⁹ were considered significant. The top-ranked genes (up to four per cluster, ranked by log fold change) were extracted for annotation. Manual annotation was performed based on (i) top-ranked DEGs and (ii) expression of canonical lineage marker genes (e.g., *EPCAM*, *CD3E*, *CD68*, *FAP*, *PECAM1*). Subclusters exhibiting mixed expression patterns of multiple major cell-type markers were defined as boundary cell populations.

### Comparison of subcluster compositions and abundances

To assess the spatial abundances of cell subclusters across treatment response groups (NR, MPR, and pCR) and timepoints (Pre, JustAfter, Resection), we used the tl.embedding_density and pl.embedding_density functions in Scanpy to visualize subcluster densities in UMAP space. For each subcluster, changes in abundance across timepoints were compared within each response category with the paired t test (e.g., Pre vs. Resection), whereas differences between response groups (e.g., NR vs. pCR) were assessed with Welch’s t test.

### Trajectory inference of CAFs using STORIES

CAFs were extracted from NR and pCR patients. To model CAF state transitions between pre-treatment biopsies (enriched for tumor-adjacent regions) and post-treatment resections (including regions farther from tumor), we focused on CAFs in close proximity to tumor cells. For each sample, CAFs located within 10 μm of tumor cells were identified using scikit-learn nearest-neighbor search (sklearn.neighbors.NearestNeighbors, n_neighbors=1) on the Xenium spatial coordinates (imagecol, imagerow) and retained for downstream analyses. For pCR resection samples in which tumor cells were absent, the tumor bed corresponding to the original tumor location was manually segmented under the supervision of a board-certified pathologist; cell IDs within the segmented tumor bed were exported from Xenium Explorer and used to include CAFs from this region. For dimensionality reduction, CAF expression matrices were scaled (sc.pp.scale) and projected to 50 principal components using Scanpy (sc.tl.pca, n_comps=50). Isomap embedding was then computed using scikit-learn (sklearn.manifold.Isomap) on the PCA representation with n_neighbors=20 and n_components=2, and the resulting two-dimensional coordinates were stored as the “isomap” basis for visualization. Trajectory inference was performed using STORIES^51^ (SpaceTime model), incorporating known temporal ordering (Pre → Resection). We fitted a stories.SpaceTime model with quadratic_weight=1e-3 using time_key="time", omics_key="X_pca", and space_key="spatial". Model optimization used AdamW with a cosine decay learning-rate schedule (optax.cosine_decay_schedule(init_value=1e-2, decay_steps=10,000) and optax.adamw). Velocities were computed from the fitted model using stories.tools.compute_velocity (with omics representation "X_pca"), and velocity fields were visualized on the Isomap embedding (basis="isomap"). All trajectory visualizations were rendered in Blender.

### Neighborhood enrichment analysis

The tumor and immune/stromal populations were combined to construct spatial neighborhood graphs using Squidpy’s gr.spatial_neighbors function. For each patient, the number of stromal, T, or myeloid cells adjacent to the tumor cells was quantified. To further assess the microenvironments surrounding the tumors, regions within 30 μm of tumor cells were extracted using the SpatialKNifeY (SKNY) algorithm^52^. Within these regions, the pairwise cell–cell adjacency was computed with Squidpy and normalized by the total edge count. Hierarchical clustering (Euclidean distance, Ward’s linkage) was used to identify cell types that frequently colocalized in tumor-adjacent microenvironments.

### Spatial stratification of Xenium data

To spatially stratify the distribution of NR CAFs, SKNY was used to extract mutually exclusive 30-μm zones surrounding tumor and normal epithelial cells. Then, the NR CAF density per unit area was calculated and compared across these regions. Stromal cells in the normal-adjacent regions, excluding NR CAFs, were redefined as fibroblasts. To define tumor cells that interacted with NR CAFs, Squidpy’s gr.spatial_neighbors was used to differentiate cells directly adjacent to NR CAFs (defined as "neighbor" cells) from those that were not adjacent to NR CAFs ("nonneighbor" cells).

### Differential expression analysis

Differential expression analysis between defined cell subpopulations was performed using the tl.rank_genes_groups function in Scanpy, in which p values and log₂ fold changes are computed according to the Wilcoxon rank-sum test.

### Ligand–receptor interaction analysis

To investigate cell–cell communication related to EMT-associated signaling pathways, we constructed an integrated AnnData object consisting of NR CAFs, pCR CAFs, fibroblasts, neighbor tumor cells, and nonneighbor tumor cells. Spatial cell–cell interaction analysis was performed using the tl.cci.grid function in the stLearn package, in which the entire spatial transcriptomics field was divided into a grid of 125 × 125 regions. A ligand–receptor database was then generated (Supplementary Table 2), focusing on interactions associated with key EMT-related pathways, including Notch signaling, TGF-β signaling, IL-6/JAK/STAT3 signaling, Wnt/β-catenin signaling, IL-2/STAT5 signaling, hypoxia, and mTORC1 signaling. We subsequently estimated the ligand–receptor interaction scores among cell types using the tl.cci.run and tl.cci.run_cci functions in stLearn.

### Visualization

Spatial localization was visualized using Xenium Explorer or the pl.spatial_scatter function from Squidpy. PhenoCycler and H&E images were visualized using QuPath. UMAP plots were generated using the pl.umap function in Scanpy.

### Survival analysis

To stratify patients on the basis of *POSTN* expression, the optimal expression cutoff value was determined with receiver operating characteristic (ROC) curve analysis, including calculation of the Youden index. Kaplan–Meier survival curves were generated with the lifelines Python library, and the statistical significance of differences was evaluated with the log-rank test.

## Supporting information

Graphical abstract

## Acknowledgments

We thank the staff of National Cancer Center Hospital East for supporting the clinical sample collections and for their helpful advice, and we thank all patients and their families for contributing to this study.

## Author Contributions

S.A.S., R.Y., and S.-I.K. conceived and designed the study. S.A.S., Y.M., K.K., M.K., Y.N., S.S., M.N., H.H., Y.T., M.I., K.S., H.B., T.K., J.Z., A.S., and Y.S. contributed to sample acquisition and data generation. S.A.S., M.O., F.H., B.R., P.V.G., and G.G. performed data analysis and interpretation. S.A.S., R.Y., and S.-I.K. wrote the manuscript. All authors reviewed, edited, and approved the final manuscript.

S.A.S., R.Y., and S.-I.K. conceived and designed the study. S.A.S., Y.M., K.K., M.K., Y.N., S.S., M.N., H.H., Y.Ts., M.I., K.S., H.B., T.K., J.Z., A.S., and Y.S. contributed to sample acquisition and data generation. S.A.S., M.O., F.H., B.R., P.V.G., and G.G. performed data analysis and interpretation. S.T., S.K., K.O., and Y.T. contributed to biological and radiation biology analyses. S.A.S., R.Y., and S.-I.K. wrote the manuscript. All authors reviewed, edited, and approved the final manuscript.

## Availability of data and materials

The clinical specimens analyzed in this study, including data generated by Xenium and Visium HD analyses, are being deposited in the DDBJ BioProject and will be made publicly available upon publication; accession numbers will be provided in the final version of the manuscript.

In accordance with collaborative research agreements and intellectual property considerations, public data release may be subject to a temporary embargo (up to 18 months) in the event of patent filing. All scripts and code used for data processing and analysis are publicly available at: https://github.com/shusakai/rectal-cancer-spatial-multiomics

## Extended Data Figures

**Supplementary Fig. 1 Marker genes of major cell clusters and interpatient variability**

**a**, Overlay of marker gene expression on a UMAP projection of the integrated Xenium dataset containing 2,829,869 cells from 52 samples across 24 patients. The genes include EPCAM and CDH1 (epithelial markers), CD3E (T cell marker), CD68 (myeloid marker), COL5A1 and COL4A1 (fibroblast/stromal markers), MYLK (smooth muscle marker), and PECAM1 (endothelial marker). **b**, Representative hematoxylin and eosin (H&E)-stained images from NR, MPR, and pCR group patients. **c**, UMAP projection of major cell clusters overlaid with patient IDs to assess interpatient variability. **d**, UMAP projection of tumor-associated major cell clusters annotated with genomic alterations in KRAS, PIK3CA, and mismatch repair (MMR) status. **e**, Violin plots showing the mean enrichment scores per patient for hypoxia and DNA repair pathways in tumor cells, stratified by pathological response. The upper panel shows Pre samples, and the lower panel shows Resection samples. For pCR cases, the Resection panel is blank because tumor cells were absent at the time of resection. Statistical significance was assessed using the Mann–Whitney U test; ns indicates no significant difference.

**Supplementary Fig. 2 Subclusters and their annotations**

**a**, UMAP plots showing subclusters of T cells, myeloid cells, and stromal cells identified by Leiden clustering. **b–d**, Dot plots showing the expression of representative differentially expressed genes (DEGs) for each subcluster identified by Leiden clustering: the top 6 genes for T cells, the top 5 genes for myeloid cells, and the top 4 genes for stromal cells. Canonical marker genes, including *EPCAM*, *CDH1*, *CD3E*, *CD8A*, *CD4*, *CD19*, *MS4A1*, *CD68*, *CD163*, *ITGAX*, *FAP*, *PDPN*, *MMP11*, *PECAM1*, *ENG*, *VCAM1*, and *MYLK*, are also shown. The dot size represents the percentage of cells expressing each gene; the color scale indicates the average expression level. **e**, UMAP visualization highlights representative clusters associated with therapy-resistant–related, response-related, and intermediate states. A dot plot shows expression of representative T cell markers and functional genes, including *CD8A*, *CD4*, *FOXP3*, *GZMA*, *CXCL13*, *TIGIT*, *PDCD1*, *CTLA4*, *LAG3*, and *IDO1*, supporting the annotation of effector CD8 T cells, regulatory T cells (Tregs), and exhausted CD8 T cells.

**Supplementary Fig. 3 Normalized abundance of cell subclusters across response groups and timepoints.**

**a**, **b**, Changes in the abundance of representative T cell, myeloid, and stromal subclusters across treatment timepoints in each response group (NR, MPR, and pCR). Line plots connect matched samples from the same patient between Pre and post-treatment timepoints. Subcluster abundance is shown as the proportion of all cells in each sample (a) or as cell number (b). P values were calculated using paired statistical tests comparing Pre and post-treatment samples. **c**, Comparison of cell density between response groups at the Pre or Resection timepoints. Each dot represents an individual sample. P values were calculated using the Mann–Whitney U test.

**Supplementary Fig. 4 Response-associated enrichment, spatial distribution, and ligand–receptor interactions of NR CAFs and NR myeloid cells**

**a**, Proportion of NR CAFs and NR myeloid cells across clinical response groups (NR, MPR, and pCR) at the resection timepoint. Each dot represents an individual sample. Statistical comparisons between groups were performed using the Mann–Whitney U test. **b**, Spatial distribution of NR CAFs and NR myeloid cells relative to tumor regions. The proportion of each cell population was calculated within distance bins from tumor cells (0–30 μm, 30–60 μm, 60–90 μm, 90–120 μm, and 120–150 μm). Lines connect measurements from the same patient. **c**, Cell–cell interaction networks among tumor cells, NR CAFs, and NR myeloid cells inferred for individual patients. Edge thickness represents the proportion of predicted interactions between cell types, and node size reflects the relative abundance of each cell population. **d**, Ligand–receptor interactions between tumor cells, NR CAFs, and NR myeloid cells. Dot plots show representative ligand–receptor pairs mediating interactions among these cell types. Color indicates the median interaction strength (MeanLR), and dot size represents the generality of each interaction across patients.

**Supplementary Fig. 5 Identification of NR CAFs and fibroblasts and analysis of the molecular function of pCR CAFs**

Spatial distribution of tumor cells (black), normal epithelial cells (gray), NR CAFs (red), and fibroblasts (blue) in patients with nonresponse (NR) (Pt-1, Pt-3, Pt-4, Pt-5, Pt-6, and Pt-8).

**Supplementary Fig. 6 Additional trajectory analyses supporting bifurcation of CAFs into NR and pCR branches.**

**a**, Flow visualization of the inferred CAF trajectory showing bifurcation into NR and pCR branches. Arrows indicate the inferred direction of cell state transitions along the manifold. Cells are colored according to branch assignment. **b,c**, Gene expression dynamics along the NR (b) and pCR (c) branches. Cells are ordered according to potential (Jθ) from Pre to Resection samples. Heatmaps show trajectory-associated genes selected by partitioning the trajectory into six stages along the potential (Jθ) and identifying the top five genes with the largest expression changes at each stage using the select_driver_genes function in STORIES.

**Supplementary Fig. 7 Identification of tumor cells interacting with NR CAFs and associated signaling pathways**

**a**, Identification and spatial localization of tumor cells adjacent to NR CAFs (neighbors) and those not adjacent to NR CAFs (nonneighbor) with the spatial neighborhood graph method in Squidpy. The plots represent the data of Pt-1, Pt-2, Pt-3, Pt-4, Pt-5, Pt-6, and Pt-8. **b**, Enrichment analysis of noncanonical WNT signaling pathways in tumor cells neighboring NR CAFs across individual patients. **c**, Predicted ligand–receptor interactions associated with NR CAF–neighbor tumor cell niches. Representative signaling pathways include angiogenesis (VEGFA–KDR), Hedgehog signaling (SHH/IHH–PTCH1), WNT signaling (WNT5A–FZD1, WNT5A–LRP6), TGF-β signaling (TGFB1/3–TGFBR1/2), TNF-α signaling via NF-κB (TNFSF10–TNFRSF10A/B and TNF–TNFRSF1A), Notch signaling (DLL1/JAG1–NOTCH1/2), and IL6/JAK/STAT3 signaling (IL6–IL6ST). Arrows indicate predicted signaling directions among NR CAFs, fibroblasts, and neighboring tumor cells.

**Supplementary Fig. 8 Graphical summary of therapy-associated CAF states and their impact on tumor**

Schematic overview summarizing the proposed model of tumor microenvironment (TME) remodeling after radiotherapy. In non-responder (NR) patients, radiotherapy induces NR CAFs adjacent to tumor cells characterized by expression of *POSTN*, *WNT5A*, and *MMP11*. These CAFs are associated with activation of multiple signaling pathways, including Notch, TGF-β, IL-6, TNF-α, and noncanonical WNT signaling, which promote epithelial–mesenchymal transition (EMT) and tumor progression, leading to a high relapse rate. In contrast, responders show the emergence of pCR CAFs in the tumor bed characterized by expression of *CXCL12*, *PI16*, and *SFRP1*, which are associated with extracellular matrix deposition and a tumor microenvironment less permissive for tumor recurrence. Normal rectal fibroblasts expressing *FGFR*, *IL10*, and *COL5A* are shown for reference.

