## Supplementary figures and images for "Single-cell spatial multiomics identifies *POSTN*^+^ CAFs mediating chemoradiotherapy resistance in rectal cancer"

### Graphical abstract

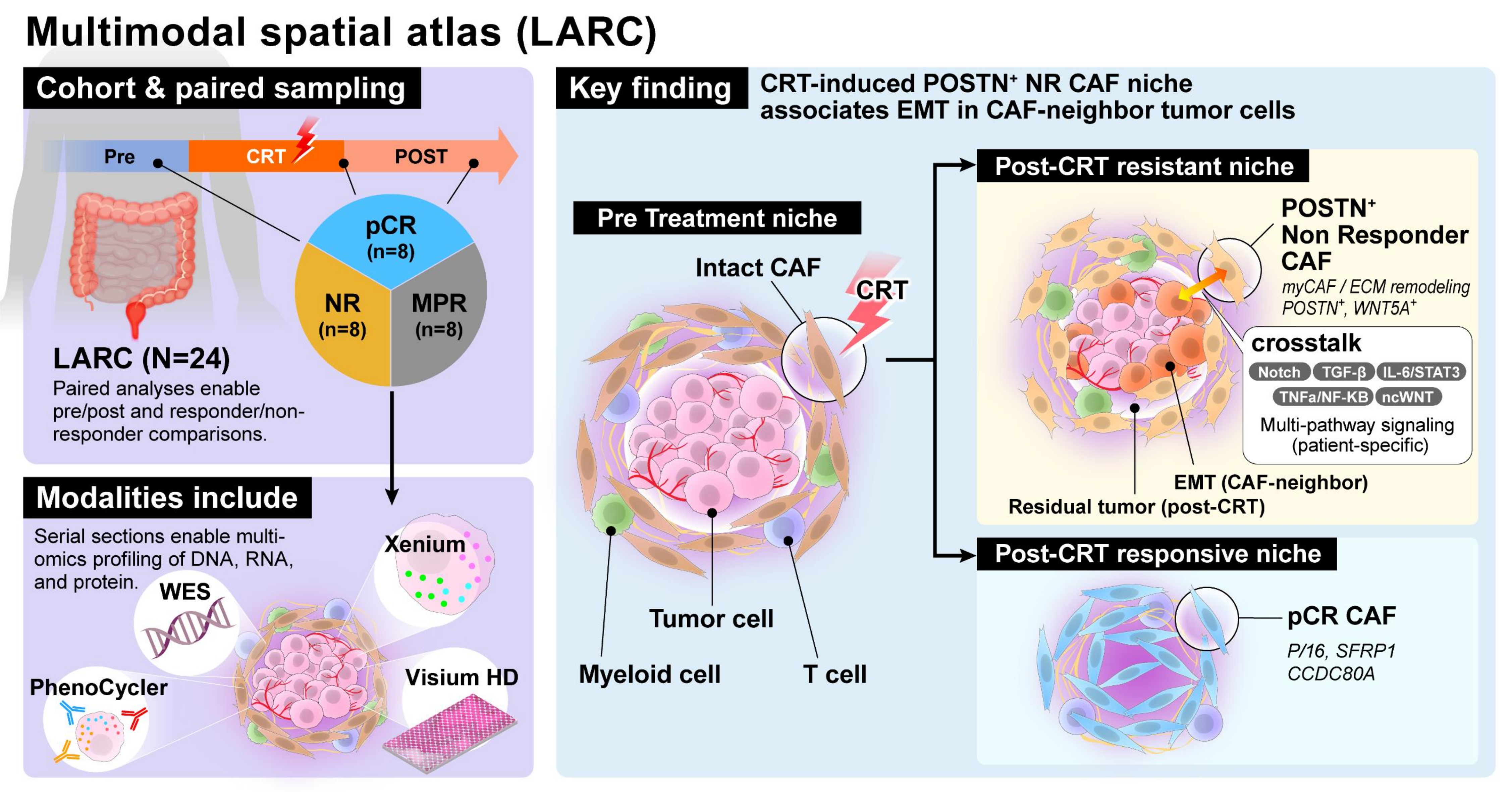

### Supplementary Fig. 1

**Supplementary Figure 1 Marker genes of major cell clusters and inter-patient variability**

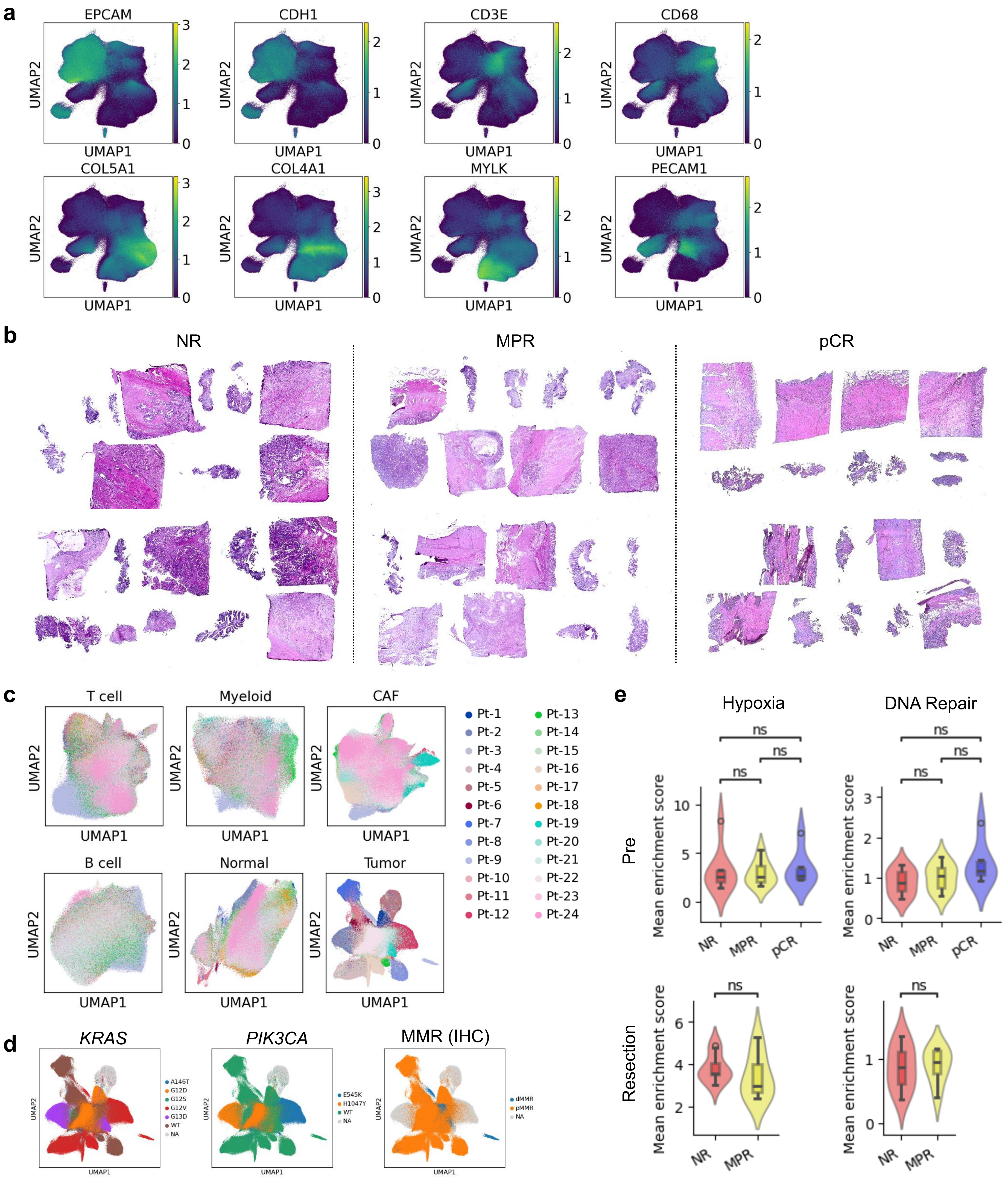

### Supplementary Fig. 2

#### Supplementary Figure 2 Subclusters and the annotations

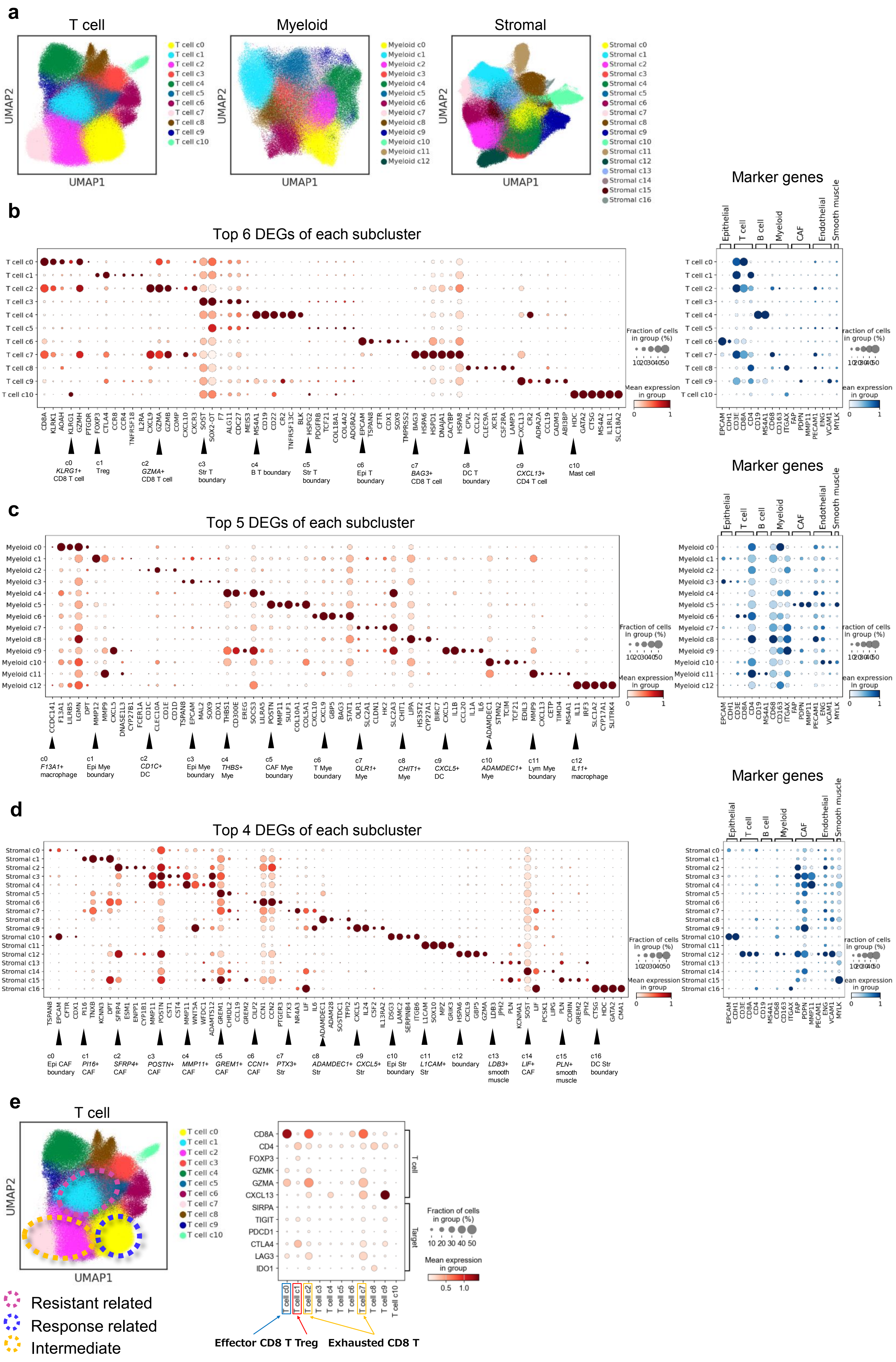

### Supplementary Fig. 3

**Supplementary Figure 3 Abundance of subclusters across response groups and timepoints.**

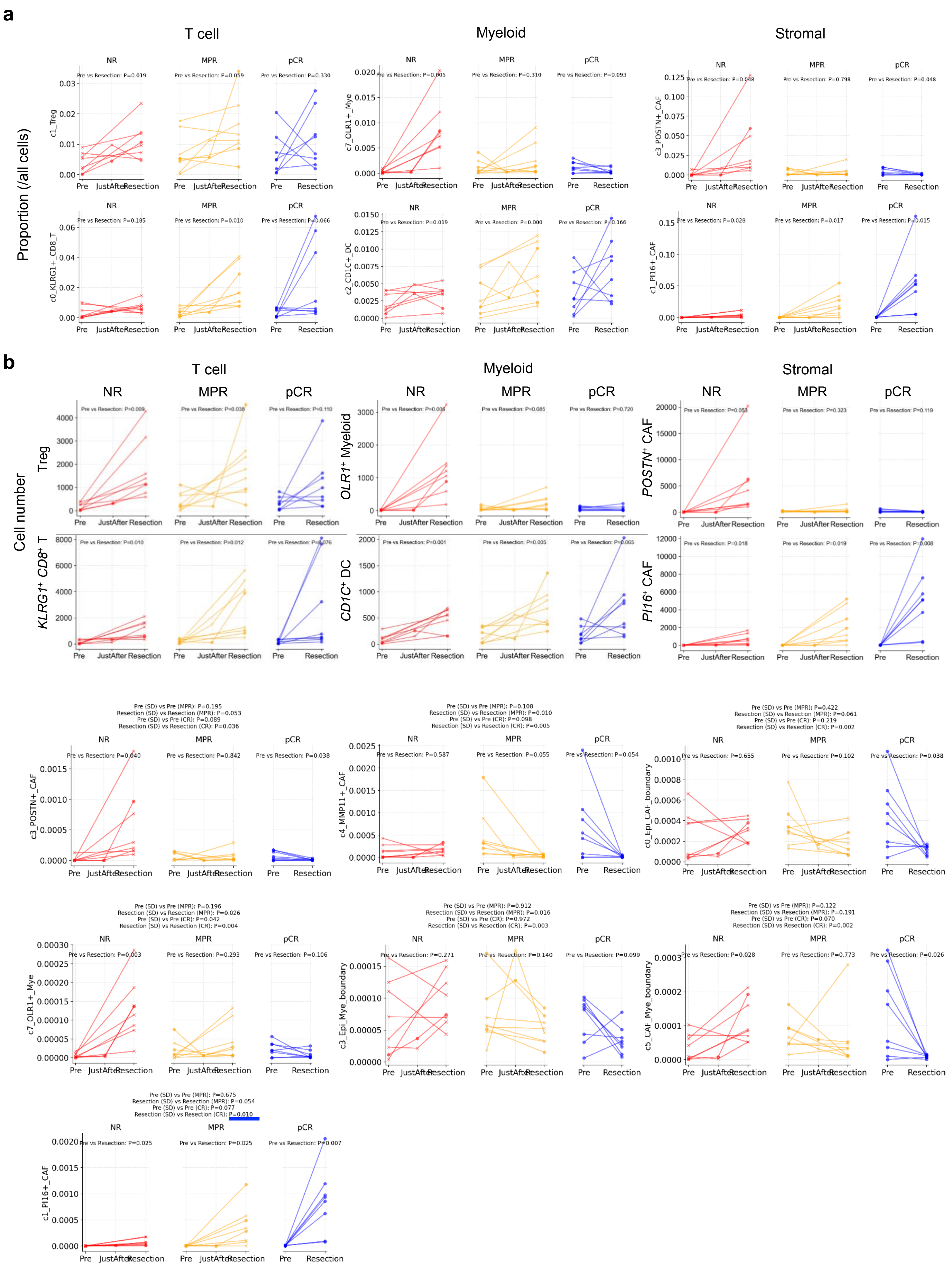
