## Supplementary Fig. 4 for "Single-cell spatial multiomics identifies *POSTN*^+^ CAFs mediating chemoradiotherapy resistance in rectal cancer"

**Supplementary Figure 4 Spatial distribution and ligand–receptor interactions among NR CAFs, NR myeloid cells, and tumor cells**

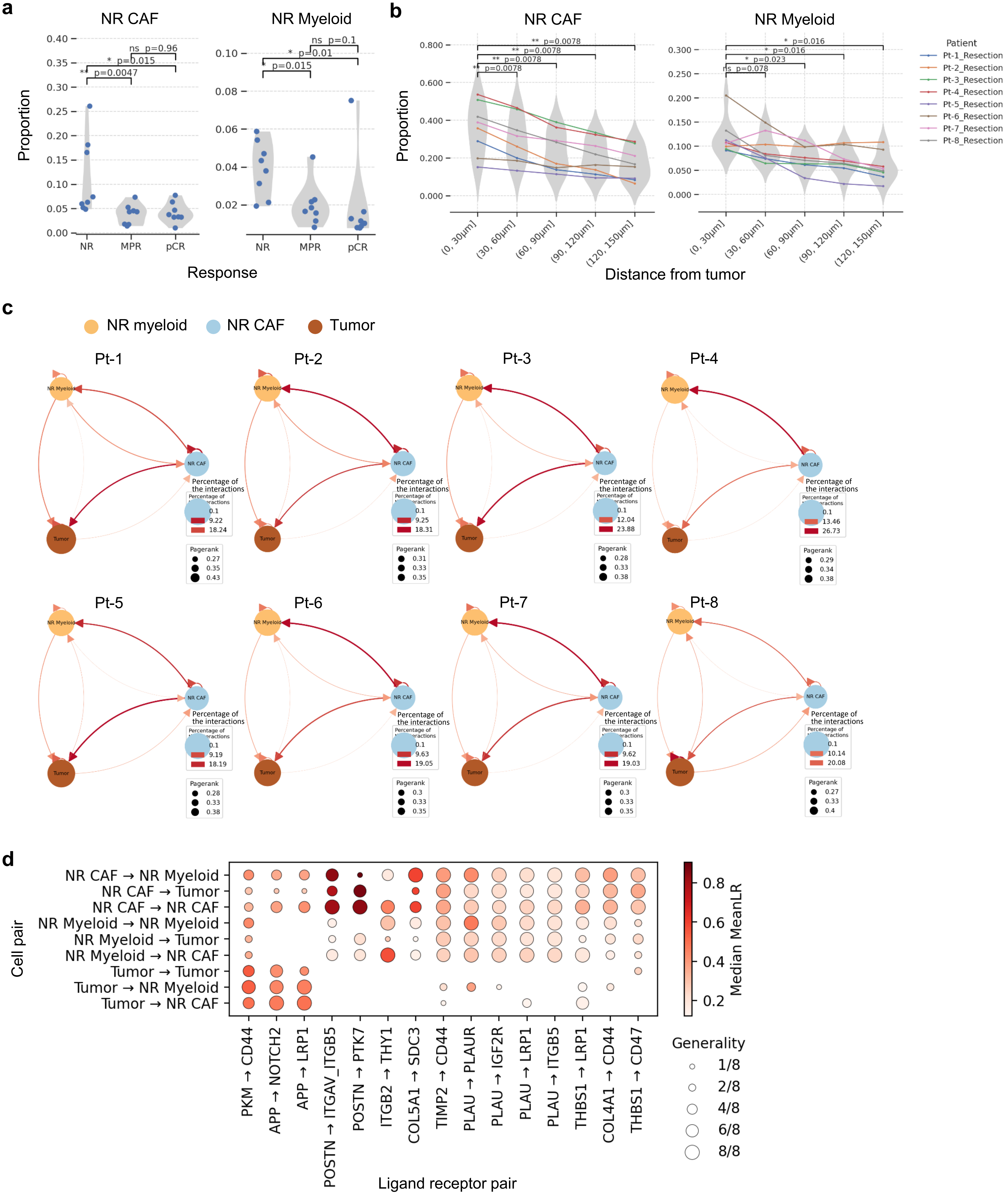
