## Supplementary Fig. 5 for "Single-cell spatial multiomics identifies *POSTN*^+^ CAFs mediating chemoradiotherapy resistance in rectal cancer"

### Supplementary Figure 5 Identification of NR CAFs and fibroblast and molecular function of pCR CAFs

**a**

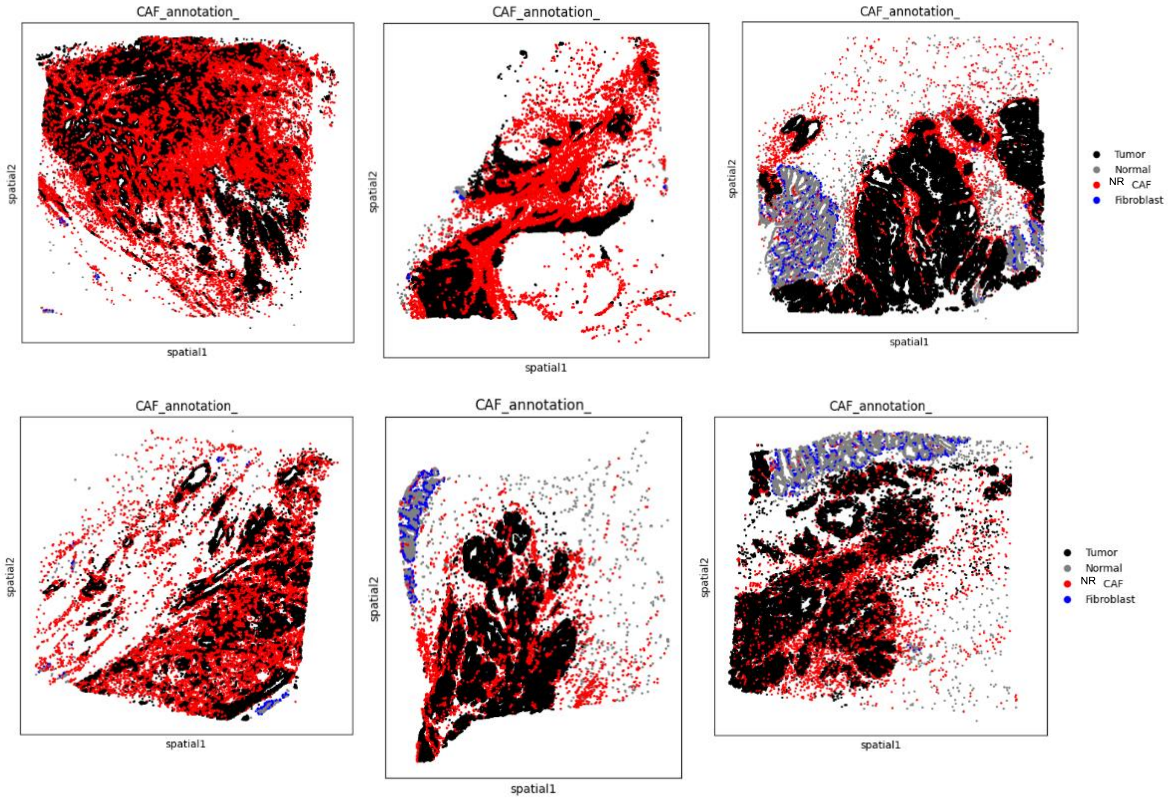
