## Supplementary Fig. 6 for "Single-cell spatial multiomics identifies *POSTN*^+^ CAFs mediating chemoradiotherapy resistance in rectal cancer"

### Supplementary Figure 6 Additional trajectory analyses of CAF bifurcation into NR and pCR branches

**a**

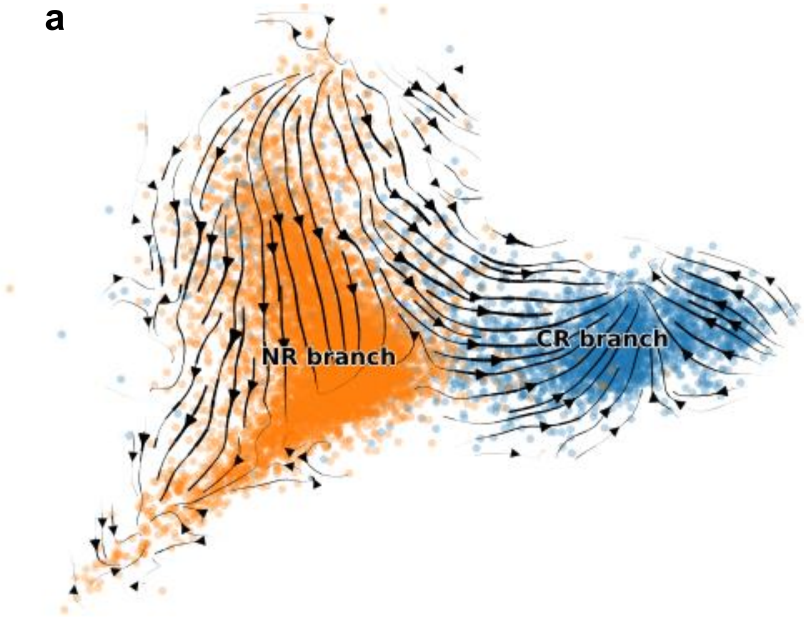


**b**

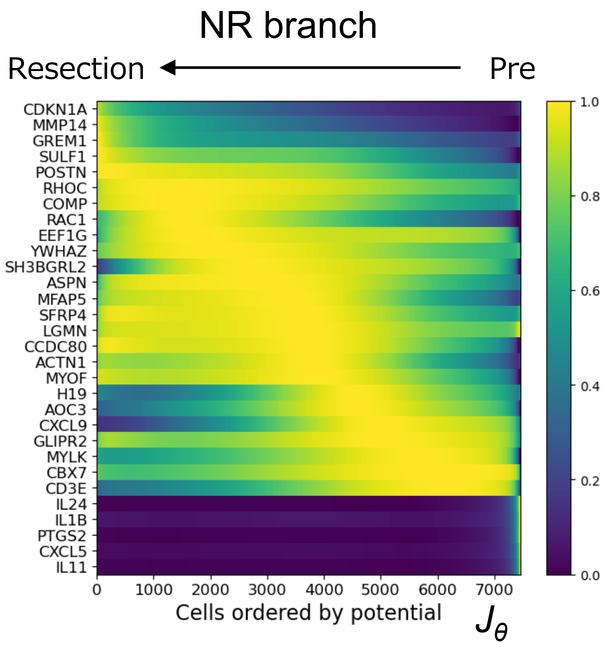


**c**

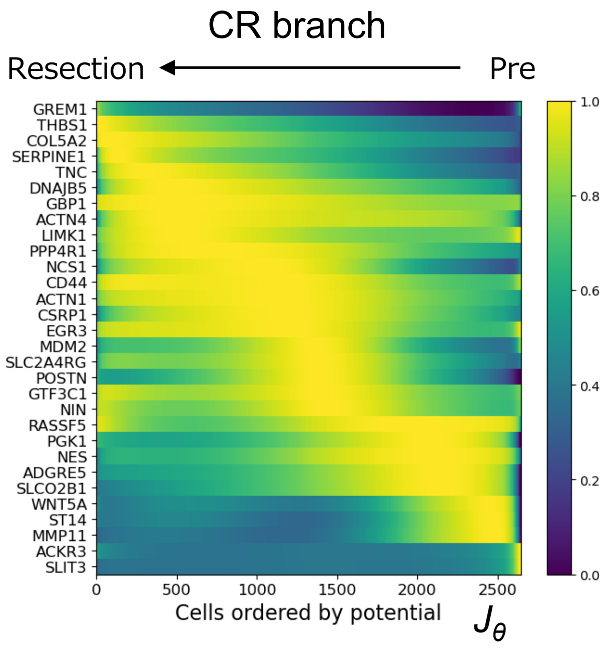
