## Supplementary Fig. 7 for "Single-cell spatial multiomics identifies *POSTN*^+^ CAFs mediating chemoradiotherapy resistance in rectal cancer"

**Supplementary Figure 7 Identification of tumor cells interacting with NR CAFs and associated signaling pathways**

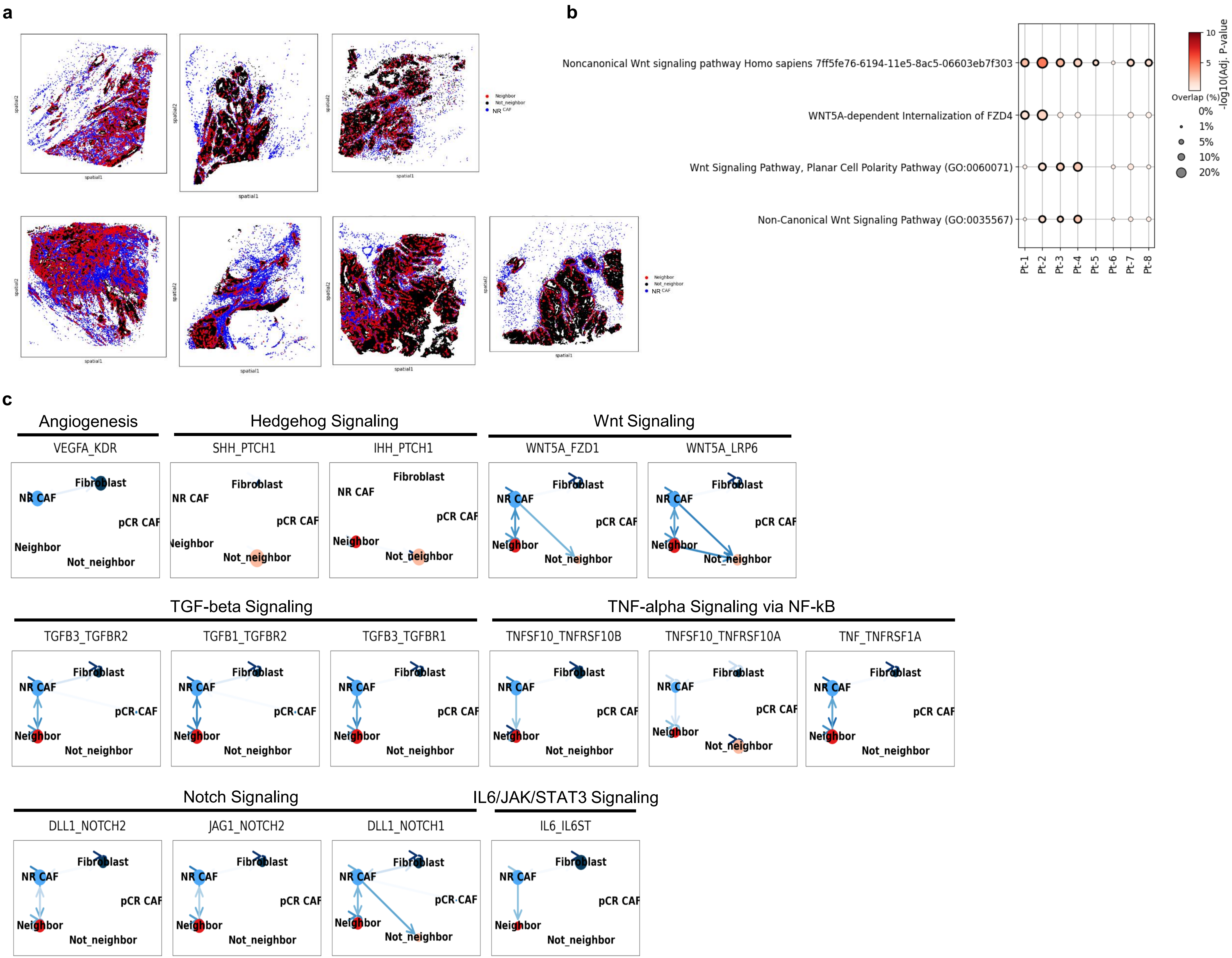
