## Supplementary Fig. 8 for "Single-cell spatial multiomics identifies *POSTN*^+^ CAFs mediating chemoradiotherapy resistance in rectal cancer"

Supplementary Figure 8 Graphical summary of therapy-associated CAF states and their impact on tumor

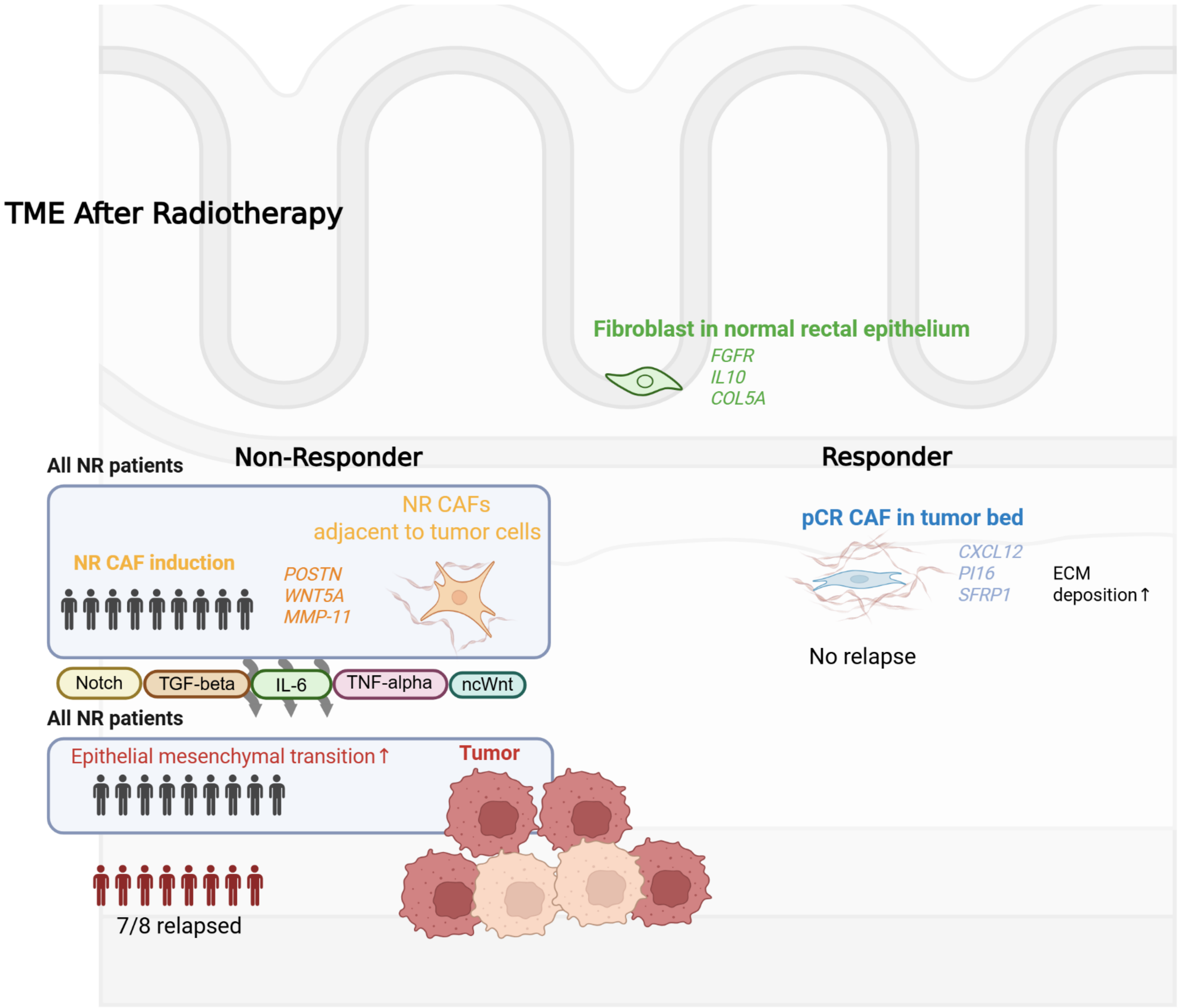
